# Arginine Regulates the Mucoid Phenotype of Hypervirulent *Klebsiella pneumoniae*

**DOI:** 10.1101/2024.11.20.624485

**Authors:** Brooke E. Ring, Grace E. Shepard, Saroj Khadka, Caitlyn L. Holmes, Michael A. Bachman, Laura A. Mike

## Abstract

Hypervirulent *Klebsiella pneumoniae* is associated with severe community-acquired infections. Hypervirulent *K. pneumoniae* colonies typically exhibit a mucoid phenotype. *K. pneumoniae* mucoidy is influenced by a complex combination of environmental factors and genetic mechanisms. Mucoidy results from altered capsular polysaccharide chain length, yet the specific environmental cues regulating this phenotype and their impact on pathogenesis remain unclear. This study demonstrates that casamino acids enhance the mucoidy phenotype but do not affect total capsular polysaccharide levels. Through targeted screening of each amino acid present in casamino acids, we identified that arginine is necessary and sufficient to stimulate the mucoid phenotype without altering capsule abundance. Furthermore, arginine activates the *rmpADC* promoter, increasing *rmpD* transcript levels, which in turn modulates capsular polysaccharide chain length and diversity. The arginine regulator, ArgR, plays a pivotal role in this regulatory cascade since deleting *argR* decreases mucoidy and increases capsular polysaccharide chain length diversity. Additionally, the Δ*argR* mutant displays increased macrophage association and has a substantial competitive defect in the lungs of mice, suggesting a link between arginine-dependent gene regulation, immune evasion and *in vivo* fitness. We discovered that arginine-dependent regulation of mucoidy is conserved in four additional hypervirulent *K. pneumoniae* isolates likely via a conserved ARG binding box present in *rmp* promoters. Our findings support a model in which arginine activates ArgR and increases mucoidy in hypervirulent *K. pneumoniae.* As a result, it is possible that arginine-dependent regulation of mucoidy allows hypervirulent *K. pneumoniae* to adapt the cell surface across different niches. This study underscores the significance of arginine as a regulatory signal in bacterial virulence.

**IMPORTANCE:** The rise of hypervirulent *Klebsiella pneumoniae* as a global health threat underscores the urgent need to understand its pathogenic mechanisms. Its ability to cause severe infections in healthy individuals and spread beyond the endemic Asia-Pacific region demands a deeper investigation into the mechanisms driving hypervirulence. The hypermucoid phenotype is primarily associated with hypervirulent isolates and is regulated by RmpD, which increases capsular polysaccharide chain length and uniformity. Understanding how environmental and genetic factors influence mucoidy is vital for elucidating the mechanisms by which *K. pneumoniae* adapts and thrives in different ecological and host niches. Our study defines the role of amino acids, particularly arginine, in regulating the bacterial surface by modulating *rmpD* expression. These results reveal that mucoidy is not a constitutive phenotype, but rather a dynamic process finely tuned by nutrient availability. Our findings expand our understanding of how the *rmp* locus is controlled and how changes in arginine availability may optimize *K. pneumoniae* immune evasion.

## INTRODUCTION

*Klebsiella pneumoniae*, a Gram-negative pathogen, ranks as the fourth deadliest bacterial species globally.^1^ Hypervirulent *K. pneumoniae* (hvKp) is a concerning pathotype that causes community-acquired outbreaks with high mortality rates ^2–5^ While hvKp infections are particularly prevalent in the Asian Pacific Rim, their incidence is spreading worldwide.^1,3,5^ This pathotype is notably dangerous due to its capacity to cause severe infections, such as pyogenic liver abscesses, endophthalmitis, and meningitis, even in healthy individuals.^2–9^ hvKp isolates are typically hypermucoid and frequently have K1 or K2 capsular serotypes.^2,3^ Hypermucoid strains sediment poorly during centrifugation and appear as sticky colonies on LB agar plates. The *rmp* locus is closely associated with hvKp strains and is found on virulence plasmids or on integrative chromosomal elements (ICEKp).^10^ This locus consists of three key components: *rmpA*, *rmpD*, and *rmpC*. In this locus *rmpA* serves as an autoregulator, *rmpC* positively regulates capsule biosynthetic genes, and *rmpD* regulates the capsular polysaccharide (CPS) chain length.^10,11^ Specifically, RmpD binding to Wzc decreases the diversity of the CPS chains on the *K. pneumoniae* cell surface.^11,12^ These changes in CPS chain length and size distribution shift the bacteria between a mucoid and non-mucoid state. Historically, capsule abundance and mucoidy were considered synonymous; however, this perception has evolved. It is now understood that this correlation is likely due to the co-expression of RmpA, RmpC, and RmpD, with RmpD being necessary and sufficient for mucoidy.^10–12^

Microenvironments within the host present different nutrient profiles to bacteria. *K. pneumoniae* mucoidy regulation is a complex process influenced by both environmental and genetic factors. Previous research has identified temperature, pH, and certain sugars (i.e., fucose) as regulating mucoidy in *K. pneumoniae*.^11,13,14^ For example, urine suppresses hvKp hypermucoidy relative to LB medium by downregulating *rmpD* transcription, while fucose increases by upregulating *rmpD* transcription mucoidy relative to glucose.^11^ However, the specific components in urine that suppress *rmpD* and hypermucoidy remain undefined. While temperature, pH, and sugar availability vary across host microenvironments, their effect on mucoidy *in situ* during infection remain unclear. In other Gram-negative bacteria, such as *Escherichia coli*, amino acid availability and distribution play essential roles in adaptation, colonization, and proliferation.^15^ However, the role of amino acids in regulating mucoidy in *K. pneumoniae* is unexplored despite multiple studies indicating that dysregulation of amino acid metabolism decreases *K. pneumoniae* pathogenesis in multiple *in vivo* models.^16,17^ Understanding how discrete host-derived nutrients modulate mucoidy during *K. pneumoniae* infections is vital for unraveling the mechanisms controlling this virulence-associated phenotype and fine-tuning bacterial fitness during pathogenesis. This is especially important given that bacterial pathogens can adapt to and exploit the metabolic resources available in their host environments to fuel their growth, regulate virulence factors, and optimize fitness.^18,19^

Amino acid availability, particularly in tissues like the urinary tract and lungs, plays a critical role in bacterial survival and pathogenesis.^15–17^ Specifically, the amino acid arginine exhibits varying concentrations across different host niches, and pathogen responses to these variations are well-documented.^15,20^ During urinary tract infections (UTIs), amino acids, including arginine, are essential for the growth and survival of uropathogenic *E. coli* (UPEC).^21^ While in the lungs, particularly during a chronic infection, arginine levels can vary significantly due to inflammation and immune response compared to that of a healthy lung. The lung becomes a rich source of metabolites, such as amino acids, when airway defenses are compromised.^22^ Specifically, in the sputum of those with cystic fibrosis, arginine is present in millimolar quantities and can serve as a key energy source for bacteria.^22,23^ Notably, pathway enrichment analysis has shown that *K. pneumoniae* arginine metabolism is upregulated during a murine lung infection.^24^

We and others have observed that culturing *K. pneumoniae* in defined media supplemented with casamino acids and glucose exhibit increased mucoidy compared to defined media with only glucose.^14^ In this study, we hypothesized that specific amino acids regulate mucoidy in *K. pneumoniae* by upregulating *rmpD* transcription and altering CPS chain length. We found that the amino acid, arginine, is necessary and sufficient to confer the hypermucoid phenotype in hvKp. This arginine-dependent regulation of hypermucoidy is dependent on the regulatory protein, ArgR. Culturing *K. pneumoniae* in arginine-containing medium results in ArgR-dependent up-regulation of *rmpD* transcription via the *rmpADC* promoter and decreased CPS chain length diversity. Arginine-dependent regulation of mucoidy is conserved in five unique hvKp strains and could be due to a predicted ARG binding box found in the promoter of the *rmp* operon. These results suggest that arginine controls mucoidy by decreasing CPS chain length diversity via ArgR-dependent activation of the *rmpADC* promoter and subsequent up-regulation of *rmpD*. Our study highlights a conserved pathway regulating one of the primary hvKp virulence factors and represents a potential anti-virulence target.

## RESULTS

### Casamino acids increase mucoidy without altering total capsule abundance

Specific nutrient sources that regulate mucoidy without affecting the total capsular polysaccharide (CPS) abundance remains elusive. To evaluate the effect of different nutrient sources on *K. pneumoniae* mucoidy and CPS, the hypervirulent strain KPPR1, was cultured in four different media: lysogeny broth (LB), low-iron minimal M9 medium supplemented with either 0.4% glucose (M9+Glc), 1% casamino acids (M9+CAA), or 0.4% glucose and 1% casamino acids (M9+Glc+CAA). The low-iron condition was used as variations in iron availability can impact CPS biosynthesis.^25^ A standard sedimentation assay was used to quantify mucoidy and a uronic acid assay was used to quantify total CPS abundance.^26^ M9+CAA significantly increased mucoidy when compared to M9+Glc (**Figure 1A**). However, M9+Glc+CAA significantly decreased mucoidy relative to M9+CAA (**Figure 1A**). Despite suppressing mucoidy, M9+Glc significantly increased CPS abundance relative to LB and M9+CAA (**Figure 1B**).

**Figure 1.**
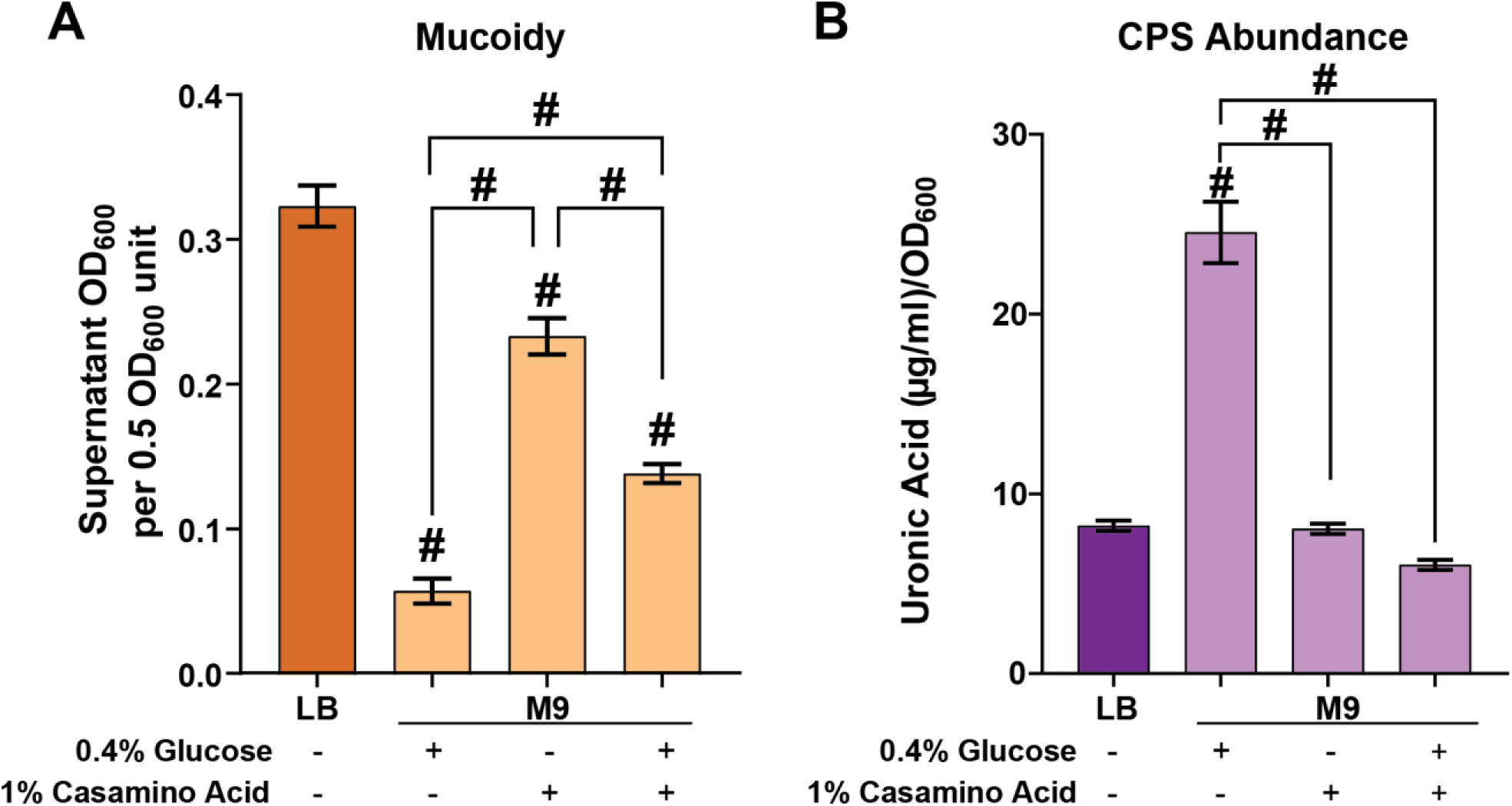
Specific nutrients dissociate CPS from mucoidy. Wild type *K. pneumoniae* strain KPPR1 was cultured in LB or low-iron M9 minimal medium with either 0.4% glucose (Glc), 1% casamino acids (CAA) or both as nutrient sources. (**A**) Mucoidy was determined by quantifying the supernatant OD_600_ after sedimenting 0.5 OD_600_ unit of culture at 1,000 x *g* for 5 min. (**B**) Uronic acid content of crude CPS extracts were quantified and normalized to the OD_600_ of the overnight culture. Data presented are the mean, and error bars represent the standard error of the mean. Statistical significance was determined using a one-way ANOVA with a Tukey post-test. The statistics displayed above the data bars represent values relative to the LB medium. The additional statistics connected by lines serve as direct comparisons to highlight differences between M9 media conditions. # p < 0.0001. Experiments were performed ≥3 independent times, in triplicate.

Furthermore, M9+Glc+CAA significantly decreased total CPS abundance compared to M9+Glc (**Figure 1B**). Together, these data indicate that culturing *K. pneumoniae* in casamino acids increases mucoidy independent of total capsule abundance.

### Arginine is necessary for the mucoid phenotype

Casamino acids is a mixture of amino acids derived from the acid hydrolysis of casein, a group of phosphoproteins present in mammalian milk. To identify which amino acid(s) in casamino acids confers mucoidy, we prepared a defined M9 medium supplemented with 20 mM sodium pyruvate and individually added 18 amino acids (M9+All) (**Supplementary Table 1**). We then generated 18 additional media, each lacking one amino acid (e.g., M9-Arg), to evaluate their individual effects on mucoidy and CPS abundance using sedimentation and uronic acid assays, respectively. We identified that compared to M9+All, the absence of three individual amino acids (arginine, phenylalanine, or glutamate) resulted in a statistically significant decrease in mucoidy without altering total CPS abundance (**Figure 2A-B**). In contrast, the absence of glycine decreased mucoidy and increased CPS abundance (**Figure 2A-B**).

**Figure 2.**
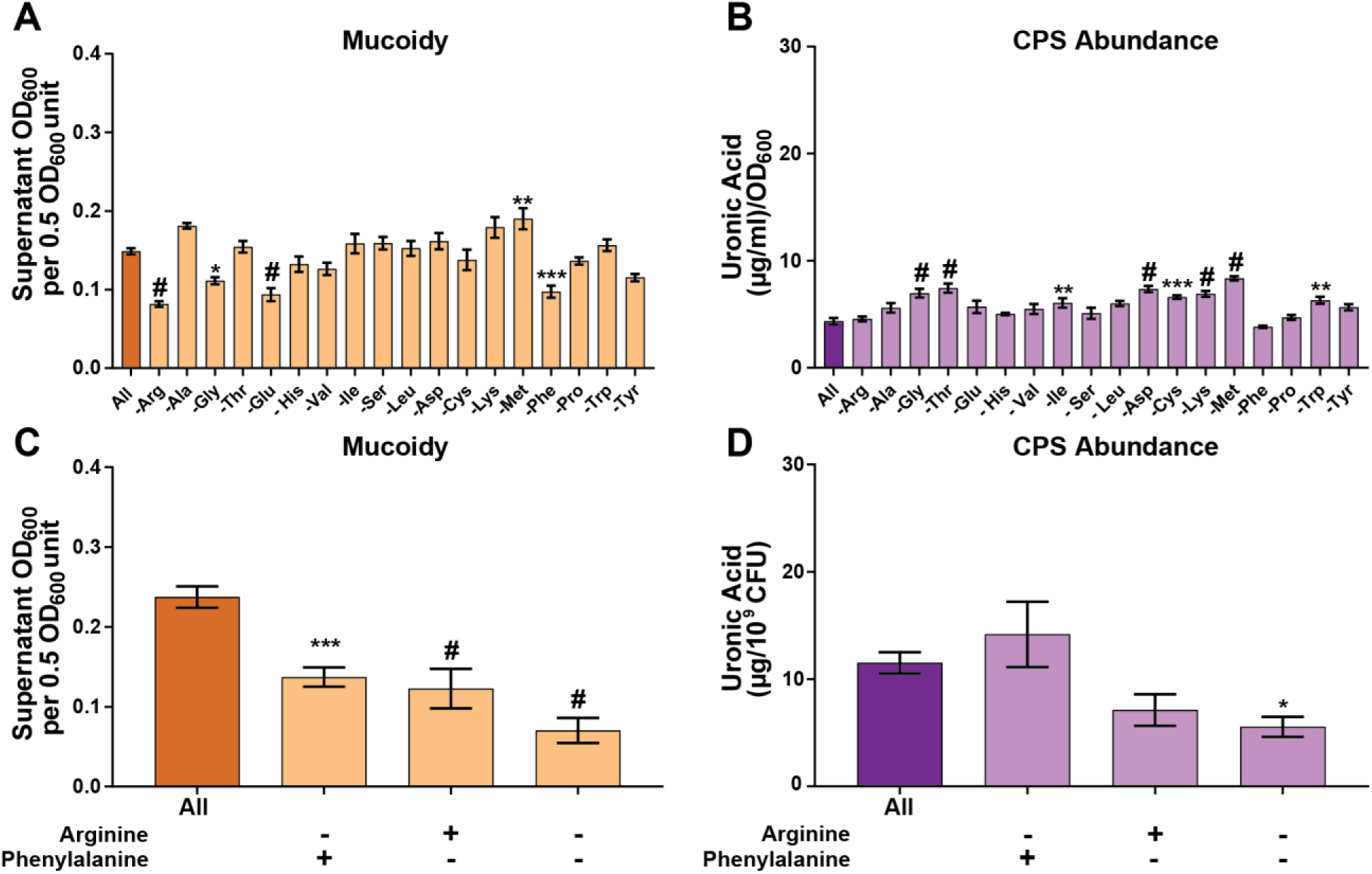
Arginine and phenylalanine are necessary for KPPR1 to regulate mucoidy. Wild type *K. pneumoniae* strain, KPPR1, was cultured in (**A, B**) low-iron M9 minimal medium with 20 mM sodium pyruvate and all 18 amino acids (M9+All) or individual amino acid absent. In (**C, D**) KPPR1 was cultured in M9+All or M9 medium without arginine, phenylalanine, or both absent. (**A, C**) Mucoidy was determined by quantifying the supernatant OD_600_ after sedimenting 0.5 OD_600_ unit of culture at 1,000 x g for 5 min. Uronic acid content of crude CPS extracts were quantified and (**B**) normalized to the OD_600_ of the overnight culture or (**D**) normalized to 10^9^ CFUs. Data presented are the mean, and error bars represent the standard error of the mean. Statistical significance was determined using a one-way ANOVA and Dunnett post-test compared to M9+All. * p < 0.05; ** p < 0.01; *** p < 0.001; # p < 0.0001. Experiments were performed ≥3 independent times, in triplicate.

Sodium pyruvate supplementation was initially included in the M9 medium to aid growth in case the absence of any amino acid impaired growth. However, we found that sodium pyruvate could suppress mucoidy when compared to M9+CAA (**Supplementary Figure 1A**). Therefore, when we validated the screening results, we prepared M9 base medium with each amino acid added at the percentage present in casamino acids but did not include 20 mM sodium pyruvate. This was critical to rule out the possibility that sodium pyruvate itself was masking or modifying the mucoid phenotype. We then prepared three M9 defined media each lacking a single amino acid identified in **Figure 2A** that decreased mucoidy (Arg, Phe, Glu, and Gly). Only the absence of arginine and phenylalanine reproduced decreased mucoidy (**Figure 2C-D**), but glutamate and glycine did not (**Supplementary Figure 1B**). We then wanted to examine if arginine and phenylalanine precursors exert the same effect and found they were not required for mucoidy (**Supplementary Figure 1B**). Ultimately, we found that only arginine absence consistently decreased mucoidy without altering total CPS abundance or growth (**Supplementary Figure 2A-C**).

### Arginine is sufficient to induce the mucoid phenotype

To identify if arginine or phenylalanine is sufficient to restore mucoidy, we screened increasing concentrations of arginine and/or phenylalanine in M9 medium, supplemented with 0.2% glycerol to aid growth (M9+Glycerol), in a 96-well plate and quantified mucoidy. We discovered that phenylalanine is not sufficient to restore mucoidy alone and that mucoidy is only restored when arginine is supplemented to the growth medium (**Supplementary Figure 3A-C)**. To validate our screen, we selected optimal arginine or phenylalanine conditions identified in the 96-well plate screen to be tested for mucoidy in our standard 3 mL culture volumes. We identified that M9 medium supplemented with 0.2% glycerol and 0.2% arginine (M9+Glycerol+Arg) is sufficient to restore mucoidy equivalent to M9+All medium, without altering capsule abundance (**Figure 3A-B**). We also assessed the growth of KPPR1 cultured in M9+Glycerol and M9+Glycerol+Arg. *K. pneumoniae* cultured in M9+Glycerol medium has an endpoint OD_600_ (0.22), which is lower than the endpoint OD_600_ of M9+Glycerol+Arg (0.44) or M9+All (0.35) (**Supplementary Figure 2,4**). Therefore, the addition of arginine can improve the growth of KPPR1. Together, these data indicate that arginine is sufficient to stimulate *K. pneumoniae* mucoidy without altering capsule abundance.

**Figure 3.**
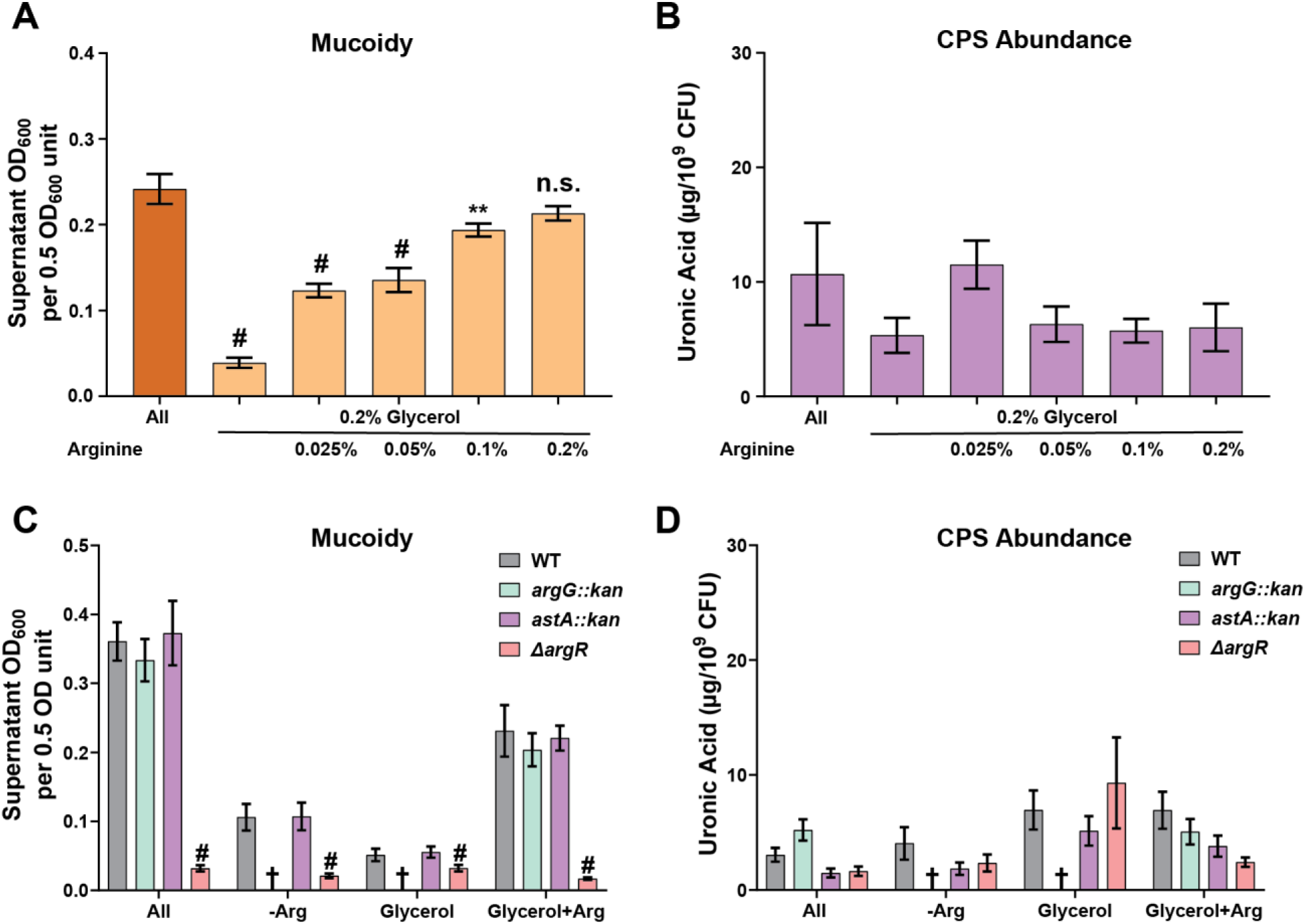
Arginine restores mucoidy in KPPR1 and ArgR is required for mucoidy regulation. **(A,B)** KPPR1 was cultured overnight in low-iron M9 medium supplemented with 18 amino acids (All), 0.2% glycerol (Glycerol), or increasing concentrations of arginine (0.025%– 0.2%) and 0.2% glycerol. Additionally, KPPR1, *astA*::kan, *argG*::kan, and Δ*argR* mutants were cultured overnight in M9 minimal media under the following conditions: with 18 amino acids (All), without arginine (-Arg), with 0.2% glycerol (Glycerol), or with 0.2% glycerol and 0.2% arginine (Glycerol+Arg). (**A,C**) Mucoidy was determined for each of the conditions by sedimenting 0.5 OD_600_ unit of culture at 1,000 x *g* for 5 min and then measuring the supernatant OD600. (**B,D**) The total capsule abundance was determined by measuring the uronic acid content of crude CPS extracts and normalized to 10^9^ CFUs. Data presented are the mean, and error bars represent the standard error of the mean. In panels (**C,D**) the cross indicates no growth in a condition. (**A,B**) Statistical significance was determined using one-way ANOVA with a Dunnett post-test. (**C,D**) Statistical significance was determined using two-way ANOVA with a Tukey post-test. Experiments were performed ≥3 independent times, in triplicate ** p < 0.01; # p < 0.0001.

### The arginine regulator, ArgR, is required for mucoidy regulation in response to arginine

To identify the pathways responding to arginine in the culture medium, we targeted genes associated with arginine degradation (*astA*), arginine biosynthesis (*argG*), and arginine regulation (*argR*). We used existing KPPR1 *argG::kan* and *astA::kan* transposon mutants and generated a KPPR1 *argR::kan* deletion (Δ*argR*).^16,27–29^ All three strains were cultured overnight in the four media conditions (M9+All, M9-Arg, M9+Glycerol, and M9+Glycerol+Arg) and mucoidy was then assessed by sedimentation assay, except *argG::kan* which cannot grow in the absence of arginine (M9-Arg, M9+Glycerol). Δ*argR* exhibited significantly reduced mucoidy compared to wild-type KPPR1 in all media conditions (**Figure 3C**). The *argG::kan* and *astA::kan* transposon mutants had no significant differences in any media conditions. The capsule abundance was not altered for any of the mutants, indicating that the effects of Δ*argR* on mucoidy are independent of total capsule abundance (**Figure 3D**). Compared to wild-type KPPR1, there were no defects in the maximum endpoint OD_600_ in any of the three mutants in M9+All, except *argG::kan* had a growth defect based on area under the curve (**Supplementary Figure 5A-C**). However, since mucoidy is measured at the 16 h endpoint and all strains have the same OD_600_ at that timepoint, the observed changes in mucoidy are likely not due to growth defects. These results indicate that arginine degradation and biosynthesis are not required to regulate mucoidy, but the ArgR regulator is required for KPPR1 to exhibit the mucoid phenotype.

### Arginine increases *rmpADC* operon promoter activity and *rmpD* transcription in an ArgR-dependent manner

To determine how arginine alters mucoidy status, we asked if increasing arginine abundance stimulates *rmpADC* promoter (P*_rmp_*) activity. To examine the effect of arginine on P*_rmp_*, we generated a reporter plasmid with DasherGFP under the control of P*_rmp_* (P*_rmp-_dasherGFP*). KPPR1 carrying P*_rmp_-dasherGFP* was cultured to stationary phase in either M9+All, M9-Arg, M9+Glycerol, or M9+Glycerol+Arg medium and then GFP fluorescence was measured and normalized to OD_600_. P*_rmp_* activity was decreased when the reporter strain was cultured in M9-Arg medium (non-mucoid) relative to M9+All medium (**Figure 4A**). Conversely, P*_rmp_* activity was increased when the reporter strain was cultured in M9+Glycerol+Arg medium (mucoid) relative to M9+Glycerol medium (**Figure 4A**). All arginine dependent changes in P*_rmp_* activation were absent in the Δ*argR* mutant (**Figure 4A**).

**Figure 4.**
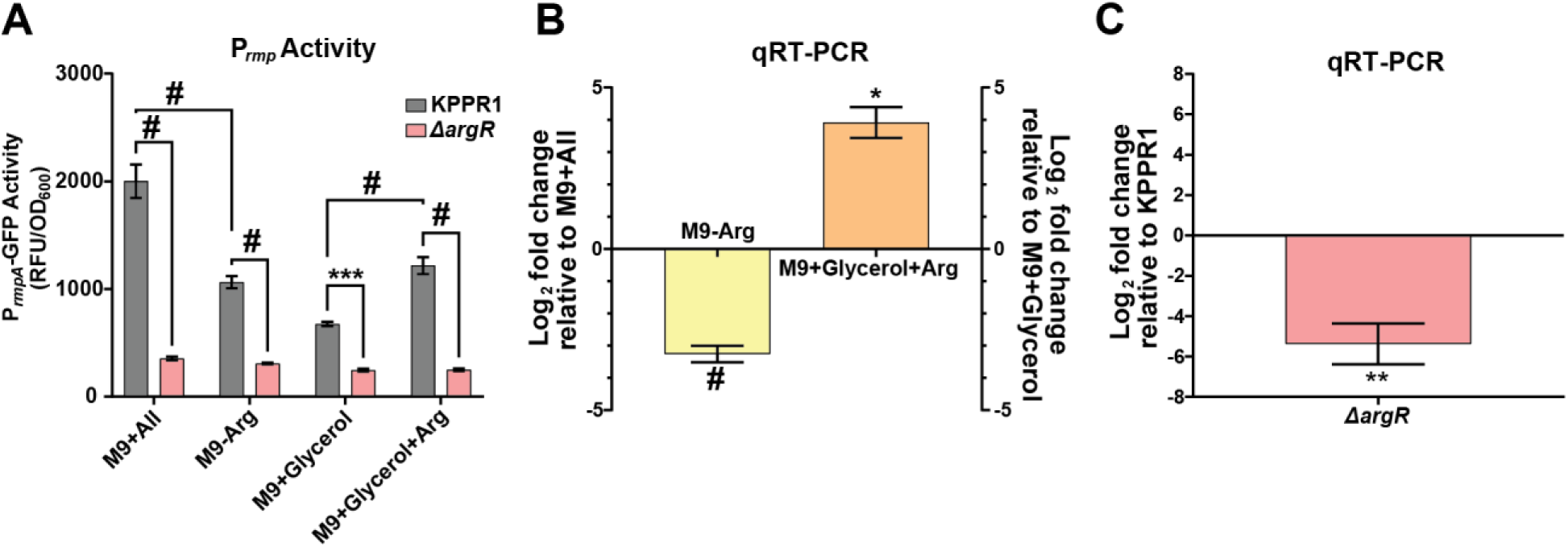
Arginine regulates mucoidy by upregulating *rmpD* transcription in an ArgR-dependent manner. KPPR1 and the Δ*argR* mutant were cultured in different amino acid conditions (M9+All, M9-Arg, M9+Glycerol, M9+Glycerol+Arg). (**A**) Promoter activity was measured by GFP expression under the control of the *rmp* operon. RNA was isolated from mid-log cultures of KPPR1 in the four media conditions or from Δ*argR* in M9+All. (**B, C**) The relative abundance of *rmpD* in the different conditions was determined by qRT-PCR and normalized to *gap2* transcript abundance. (**B**) Relative *rmpD* transcript levels in KPPR1 cultured in M9-Arg or M9+Glycerol+Arg were compared to M9+All or M9+Glycerol, respectively, while (**C**) the Δ*argR* mutant cultured in M9+All transcript levels were compared to wildtype, KPPR1. Data presented are the mean, and error bars represent the standard error of the mean. Statistical significance was determined in panel **A** with a two-way ANOVA with Tukey’s post-test to compare wild-type KPPR1 to the Δ*argR* mutant and comparing wildtype-KPPR1 in the four conditions. In panels **B** and **C** significance was determined using a student’s t-test to determine if either group were significantly different than 1. * p<0.05; ** p < 0.01; *** p < 0.001; # p < 0.0001. Experiments were performed ≥3 independent times, in triplicate.

As we and others have shown, *rmpD* transcript levels correlate with mucoidy levels due to RmpD modulating Wzc function to increase CPS chain length and uniformity.^11,12^ To examine the effects of the absence and presence of arginine on *rmpD* expression, we quantified *rmpD* and *gap2* (internal control) transcript abundance in KPPR1 cultured in the four same media conditions (M9+All, M9-Arg, M9+Glycerol, and M9+Glycerol+Arg). In M9-Arg medium (non-mucoid), the *rmpD* gene is significantly downregulated compared to M9+All medium (**Figure 4B**). Additionally, in M9+Glycerol+Arg medium (mucoid), *rmpD* transcripts are significantly upregulated compared to M9+Glycerol (**Figure 4B**). We found a significant decrease in *rmpD* transcripts in the Δ*argR* mutant strain relative to wild-type KPPR1 (**Figure 4C**). No other candidate amino acids or genes changed *rmpD* transcript levels (**Supplementary Figure 5F; Supplementary Figure 6A**). These results indicate that the presence of arginine increases mucoidy by increasing P*_rmp_* activity and *rmpD* transcription in an ArgR-dependent manner.

### Arginine and ArgR decrease diversity of capsular polysaccharide chains on the cell surface

Recently, our lab and others reported that diverse, shorter CPS chains are associated with a non-mucoid colony phenotype while uniform, longer CPS chains are associated with a hypermucoid phenotype.^11,12^ RmpD interacts with the chain length regulating auto-kinase, Wzc, to increase CPS chain length and uniformity.^11,12^ Therefore, we examined the effects of arginine and ArgR on CPS chain length diversity. Cell-associated CPS were isolated after wild-type KPPR1 was cultured in either M9+All, M9-Arg, M9+Glycerol, or M9+Glycerol+Arg.^26^ The cell-associated CPS purified from 1.5 OD_600_ of bacteria and resolved by SDS-PAGE and visualized using Alcian blue and then silver stain.^26^ When KPPR1 is cultured in LB medium the bacteria appear mucoid and produce three distinct polysaccharide bands, corresponding to CPS species that we previously described as ’Type A’ (diverse; mid-to high-molecular weight chains), ’Type B’ (uniform; high-molecular weight chains), and ’Type C’ (ultrahigh-molecular weight chains).^11^ When arginine is absent from the media, the diverse Type A chains that appear as a smear on the gel increase significantly, and when arginine is present in the media, the diverse Type A chains decrease significantly (**Figure 5A-B**). We evaluated the CPS chain length diversity of wild type and Δ*argR* cultured in M9+All medium, as we observed the strongest mucoid phenotype differences in this culture medium (**Figure 3C**). Δ*argR* produced significantly more ’Type A’ CPS chains and trended towards decreased ‘Type B’, thereby increasing overall cell surface CPS chain length diversity compared to wild-type KPPR1 (**Figure 5A, 5C**). No other candidate amino acids or genes altered capsule chain length diversity (**Supplementary Figure 5D-E; Supplementary Figure 6B-C**). These results indicate that the presence of arginine decreases Type A CPS abundance, increasing the overall uniformity of the cell-associated CPS chains and increasing the mucoid phenotype in an ArgR-dependent manner.

**Figure 5.**
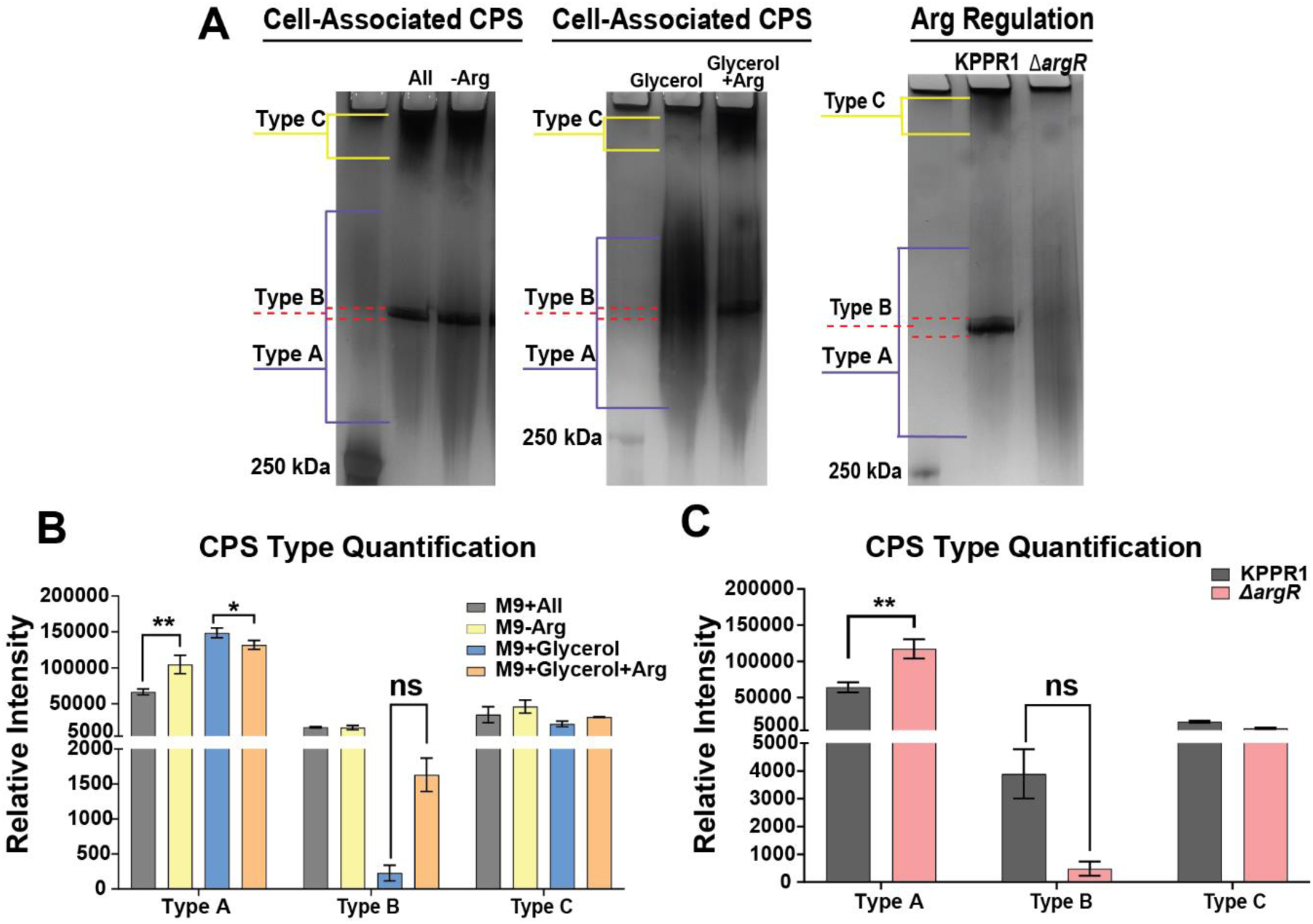
Arginine and ArgR regulate mucoidy by upregulating *rmpD* transcription and decreasing capsule Chain diversity. KPPR1 and the Δ*argR* mutant were cultured in different amino acid conditions (M9+All, M9-Arg, M9+Glycerol, M9+Glycerol+Arg) or (M9+All), respectively. (**A**) Capsular polysaccharides (CPS) were separated on a 4-15% SDS-PAGE gel and stained with alcian blue and silver stain. Three distinct polysaccharide types emerged: diverse mid-to high-molecular-weight chains (Type A), uniform high-molecular-weight chains (Type B), and diffuse ultra-high-molecular-weight chains (Type C). Data presented are the mean, and error bars represent the standard error of the mean. Representative images are shown in panel **A** with the quantification being shown in panels **B** and **C**. Statistical significance was determined in panel **B** using a two-way ANOVA with Tukey’s post-test was used to compare specific groups, while panel **H** a one-way ANOVA with a Dunnett’s post-test was used. * p < 0.05; ** p < 0.01. Experiments were performed ≥3 independent times, in triplicate.

We previously reported that Wzc phospho-status is sometimes reduced by Wzc mutations that increase mucoidy and Types B and C polysaccharide production.^11^ Therefore, we also determined whether the absence of arginine or phenylalanine affects Wzc phospho-status.^11^ KPPR1 was cultured in M9+All, M9-Arg, M9-Phe, and M9-Arg-Phe and whole-cell lysates were prepared and separated by SDS-PAGE. The phospho-tyrosine profiles in each media condition were analyzed by Western blot and we found Wzc phospho-status is only decreased when both arginine and phenylalanine are absent (**Supplementary Figure 6D-E**). Although the literature supports that Wzc phosphocycling directs CPS chain length regulation, our findings suggest that Wzc phospho-status is unlikely to be a good indicator of CPS chain length diversity regulation. It is possible that monitoring phospho-kinetics may provide more insight into the regulation of the diversity of CPS chains. Nonetheless, our data indicate that arginine and ArgR decreases CPS chain length diversity, likely through RmpD-Wzc interactions.

### ArgR reduces macrophage association and enhances bacterial fitness in the lung

To examine the role of ArgR in host interactions, we validated that Δ*argR* is responsible for regulating mucoidy via complementation. We restored mucoidy to Δ*argR* by expressing the *argR* gene under the control of the *argR* promoter (P*_argR_*) *in trans* (**Figure 6A**). Additionally, we restored mucoidy to Δ*argR* by expressing the *rmpD* gene under the control of the *rmpD* promoter (P*_rmpD_*) *in trans* (**Figure 6A**). This indicates that ArgR acts above RmpD in the pathway regulating mucoidy. After complementation, we then quantified how effectively Δ*argR* adheres to immortalized bone marrow-derived macrophages from *Mus musculus* (iBMDMs, BEI #NR-9456).^11^ We found the Δ*argR* mutant bound macrophages 3.2-fold more than WT (P < 0.00001) (**Figure 6B**). To determine the effects of an *argR* deletion during infection, we competed WT_EV_ against Δ*argR*_EV_ or Δ*argR*_P*argR*_ in a murine pneumonia model. As predicted, the Δ*argR* mutant carrying the empty plasmid vector was out competed 2.5-fold by WT_EV_ (P < 0.0001) (**Figure 6C**). While there was a trend towards reduced fitness of Δ*argR*_EV_ in the spleen and liver, there was not a significant difference between Δ*argR*_P*argR*_ and WT_EV_. Taken together these data indicate that ArgR activity blocks bacterial adherence to and internalization by host immune cells and is necessary for bacterial fitness in the lungs.

**Figure 6.**
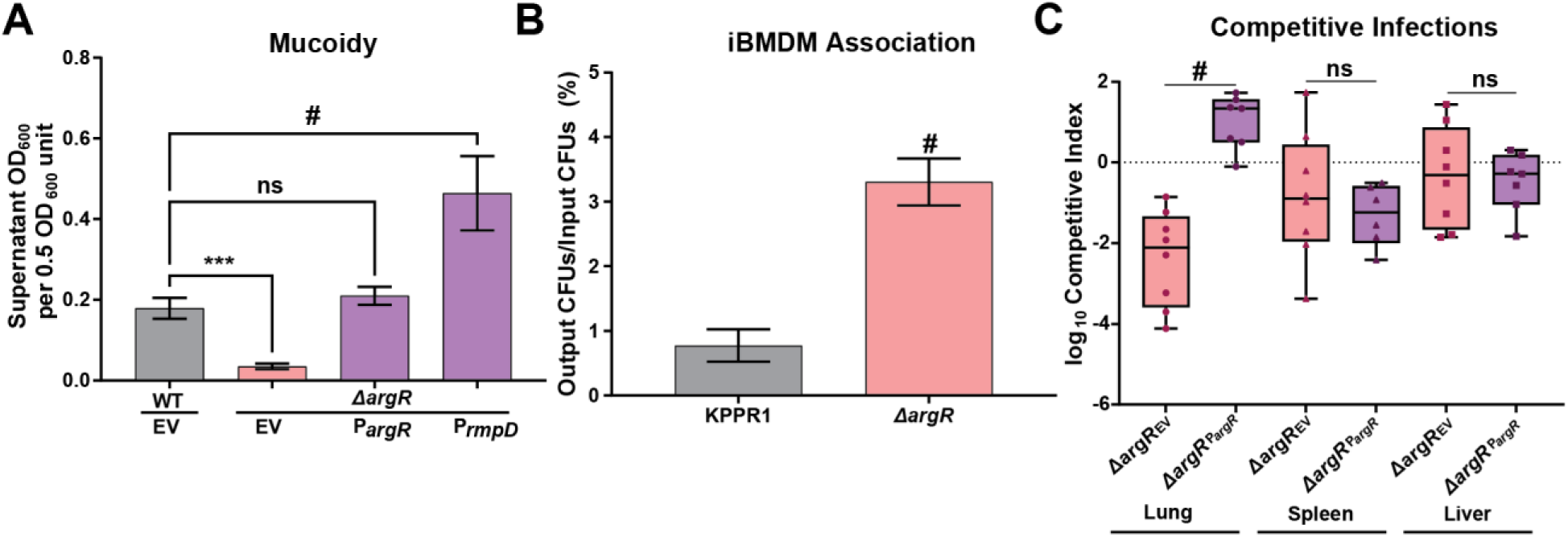
ArgR blocks bacterial association with immortalized bone marrow-derived macrophages (iBMDMs) and promotes competition fitness in the lung. The strains KPPR1 and Δ*argR* were transformed with either the empty vector (EV) or *argR* (P*argR*) or *rmpD* (P*rmpD*). The EV backbone is pACYC184Δ*tet*. (**A**) Wildtype and the Δ*argR* mutant carrying the vectors were cultured overnight in M9+All and mucoidy was determined by a sedimentation assay (0.5 OD_600_ unit centrifuged at 1,000 x *g* for 5 min). Macrophages (BEI NR-9456) were incubated with KPPR1 wildtype or Δ*argR* for 2 hours (MOI = 10). (**B**) Macrophages were washed with PBS 3x, then lysed with 0.2% TritonX100. CFUs were enumerated and normalized to input CFUs for association. Data presented (**A**, **B**) are the mean, and error bars represent the standard error of the mean. (**C**) KPPR1 EV and Δ*argR* carrying EV or P*argR* were cultured overnight and 1x10^6^ CFU of 1:1 mixture of WT_EV_ and Δ*argR*_EV_ or Δ*argR*_P*argR*_ .were administered retropharyngeally to C57BL/6 mice. The log_10_ competitive index at 24 h post infection is shown relative to 1.0 for individual mice with bars representing the median and interquartile range. Statistical significance was determined in panel A with a one-way ANOVA and Dunnett’s post-test, while in panel **B** a paired t-test was used. In panel **C,** an unpaired t-test was used to determine statistical significance. *** p < 0.001; # *p* < 0.0001. Experiments were performed ≥2 independent times, with each dot representing an individual mouse

### Arginine-dependent regulation of mucoidy is consistent across multiple hypervirulent *K. pneumoniae* isolates

As the data thus far were collected using the hvKp strain KPPR1, we next evaluated if arginine-dependent control of mucoidy is conserved in other hvKp strains. We selected the laboratory strain, NTUH-K2044, and three hvKp clinical isolates (Kp4289, Kp4585, and Kp6557) for further study.^30^ First, we used PathogenWatch to confirm that all strains were *rmp-*positive *K. pneumoniae* (**Table 1**).^31–37^ To evaluate if mucoidy was up-regulated by arginine in these four hypervirulent strains, the strains were cultured in M9+All or M9-Arg and mucoidy was quantified by sedimentation assay. We found that all strains had a significant increase in mucoidy when cultured in M9+All relative to arginine-free medium (M9-Arg) (**Figure 7A**). These results indicate that the positive regulation of mucoidy by arginine occurs in other hvKp strains.

**Figure 7.**
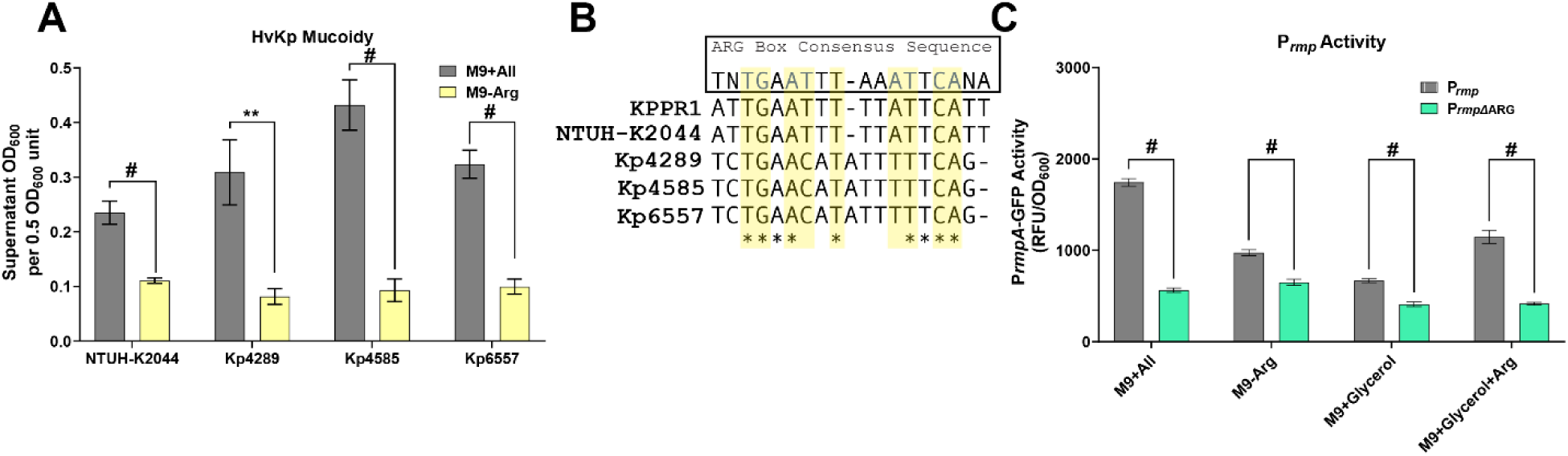
Multiple HvKp strains regulate mucoidy in response to arginine and encode an ARG binding box in the *rmp* promoter. One hypervirulent *K. pneumoniae (*hvKp) laboratory strain (NTUH-K2044) and three hvKp clinical isolates (Kp4289, Kp4585, and Kp6557) were cultured in M9+All or M9-Arg. (**A**) Mucoidy was assessed by quantifying the supernatant OD_600_ after sedimenting 0.5 OD_600_ unit of culture at 1,000 x *g* for 5 min. (**B**) A BPROM-predicted ARG binding box from the *rmp* promoter of five HvKp strains was aligned with ClustalW. The yellow highlighted nucleotides represent highly conserved nucleotides in the ARG box with stars representing conservation of the nucleotide across all five strains. (**C**). KPPR1 carrying either the P*_rmp_*-GFP reporter plasmid or a scrambled ARG box (P*_rmpΔ_*_ARG_-GFP) was cultured in different amino acid conditions (M9+All, M9-Arg, M9+Glycerol, M9+Glycerol+Arg). Promoter activity was measured by GFP expression under the control of the *rmp* operon. Data presented are the mean, and error bars represent the standard error of the mean. Statistical significance was determined using an unpaired t-test. ** p < 0.01; # p < 0.0001. Experiments were performed >3 independent times, in triplicate.

### An ARG binding box in the *rmp* promoter regulates activity in response to arginine

ArgR is activated by arginine and binds to regulatory targets via an ARG binding box. Therefore, we used BPROM, a bioinformatic bacterial promoter prediction tool, to identify a predicted ARG box in the *rmp* promoter of KPPR1 and four additional hypervirulent strains.^38^ Using CLUSTALW, we aligned the five predicted ARG boxes and generated a consensus sequence (**Figure 7B**). The ARG box in *K. pneumoniae* includes eight nucleotides that are highly conserved across all ARG boxes, with additional nucleotides that may vary. (**Figure 7B**, highlighted yellow).^39^ Seven out of the eight conserved ARG box nucleotides were present in the three clinical isolates, while all eight ARG box nucleotides were identified in KPPR1 and NTUH-K2044 (**Figure 7B**).^39^ To determine if the ARG box is required for arginine-dependent regulation, we mutated the eight conserved nucleotides in P*_rmp_* on the reporter plasmid to make P*_rmp_*_ΔARG_-dasherGFP. WT KPPR1 carrying P*_rmp_*_ΔARG_ and P*_rmp_*_ΔARG_ reporter plasmids were cultured in either M9+All, M9-Arg, M9+Glycerol, or M9+Glycerol+Arg medium and measured GFP fluorescence as before. GFP expression was decreased in KPPR1 carrying the P*_rmp_*_ΔARG_ in all media compared to KPPR1 P*_rmp_* (**Figure 7C**). These results suggest that ArgR regulates mucoidy (**Figure 3A**) in response to arginine availability (**Figure 2C**) by binding to the ARG binding box in the promoter of *rmpADC* (**Figure 7B-C**), increasing expression of the *rmpADC* operon (**Figures 4A-C**), producing less diverse CPS chains (**Figures 5A-C**), decreasing association with macrophages and fitness in the lung (**Figure 6B-C**), and increasing mucoidy in hypervirulent *K. pneumoniae* strains (**Figures 7A** and **8**).

**Figure 8.**
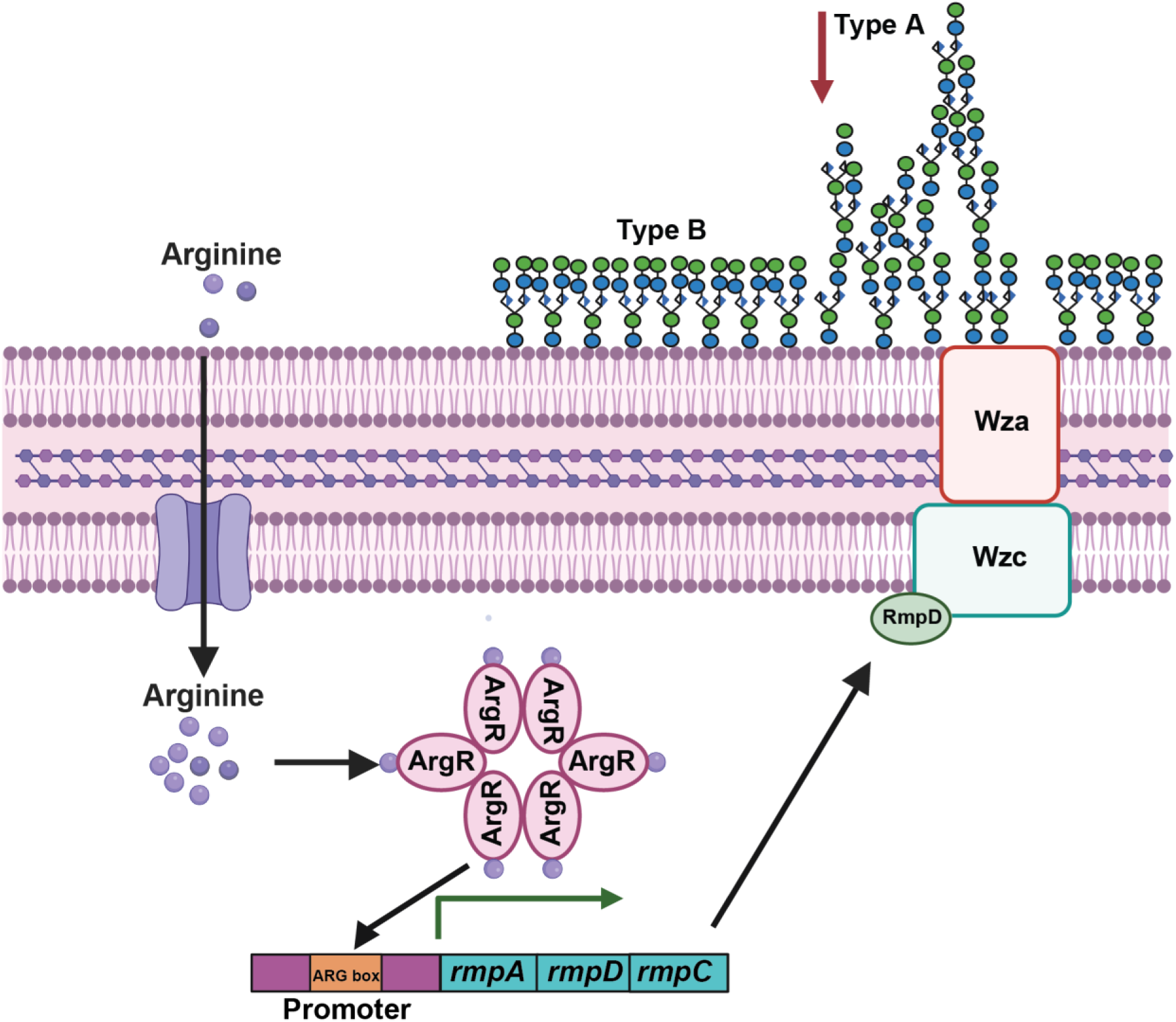
Model of *K. pneumoniae* regulation of mucoidy in response to arginine. When cultured in the presence of the semi-essential amino acid, arginine, *K. pneumoniae* increases mucoidy without altering the total capsule abundance. When arginine is brought into the cell, the arginine regulator, ArgR, will bind with arginine and become active. The activated ArgR-arginine can act as a transcriptional regulator by binding to ARG boxes. The arginine-ArgR complex activates the *rmpADC* promoter in an ARG-box dependent manner and upregulates *rmpD*, the small protein known to regulate mucoidy. The RmpD protein then goes on to interact with the Wzc protein, a tyrosine kinase that regulates high-level capsule polymerization. RmpD-Wzc interactions decrease ‘Type A’ and increase ‘Type B’ polysaccharide chains, which decreases CPS diversity and presents as the mucoid phenotype. *Created with Biorender.com*

## DISCUSSION

*K. pneumoniae* mucoidy is influenced by both environmental conditions and genetic factors, with arginine emerging as a key modulator of this phenotype. Our study reveals that arginine regulates mucoidy by activating the virulence-associated *rmp* locus, which in turn alters capsule properties. Our findings reveal that while casamino acid significantly increase mucoidy, they do not alter the overall total capsular polysaccharide abundance (**Figure 1A-B**). The distinct effect of casamino acids on mucoidy and not capsule abundance is a critical observation as it aligns with recent work demonstrating that capsule abundance and mucoidy are distinct from one another.^11,12^ Mucoidy is now attributed to reduced CPS chain length diversity rather than simply to higher capsule abundance. While this manuscript focuses on how amino acids regulates mucoidy, we did observe that glucose suppresses mucoidy in the presence of casamino acids, indicating a complex interaction between different nutrients (**Figure 1B**).

A systematic analysis of the contribution of each amino acid revealed that arginine is the key amino acid driving mucoidy (**Figure 2-3**). Subsequent validation confirmed that the absence of arginine decreased mucoidy without affecting capsule abundance or growth (**Figure 2** and **Supplemental Figure 1**). These data demonstrated that arginine is necessary for the regulation of *K. pneumoniae* mucoidy. Moreover, when arginine is supplemented into culture medium with glycerol as the sole carbon source, mucoidy is restored to the same level observed in M9 with all amino acids (M9+All) without altering capsule abundance (**Figure 3A-3B**). These data demonstrated that arginine is sufficient to positively regulate mucoidy independently of capsule abundance.

To determine how arginine regulates *K. pneumoniae* mucoidy, we investigated if genes involved in arginine biosynthesis, degradation, or regulation regulate mucoidy. We found that the arginine regulator, ArgR, is required to stimulate *K. pneumoniae* mucoidy (**Figure 3C-D**). ArgR is known to coordinate arginine synthesis and transport, but can also regulate other genes.^40–42^ Previous studies have shown that in hypervirulent *K. pneumoniae* strain NTUH-K2044, the Δ*argR* mutant exhibits decreased mucoidy and capsule abundance with slightly reduced capsule thickness and finer filaments as visualized by electron microscopy.^40^ Our KPPR1 Δ*argR* mutant mirrors the reduced mucoidy of NTUH-K2044 Δ*argR*, strengthening our conclusion that ArgR is a key mucoidy regulator (**Figure 3C**).

Previous work has shown that RmpD interacts with Wzc to reduce CPS chain length diversity and produce the mucoid phenotype.^12^ Here, we demonstrate that arginine activates the *rmpADC* promoter, increases *rmpD* transcription and influences CPS chain length (**Figure 4A-C**).^10–12^ Specifically, the absence of arginine, which results in the loss mucoidy, increases CPS chain length diversity. This effect on CPS chain length diversity is reversed when arginine is reintroduced, suggesting that arginine plays a role in reducing capsule chain diversity to enhance mucoidy (**Figure 5A-B**). Our data support that the effects of arginine on CPS chain length are likely mediated by ArgR, as ArgR is required for increasing *rmp* promoter activity and *rmpD* transcription in the presence of arginine (**Figures 4A**). Δ*argR* also exhibits increased CPS chain length diversity compared to WT (**Figure 5A, 5C**).

The role of ArgR in regulating mucoidy via the *rmp* promoter was validated by restoring mucoidy by complementing Δ*argR* with either *argR* or *rmpD in trans* (**Figure 6A**). Restoring mucoidy to Δ*argR* by expressing *argR* under its own promoter confirmed that *argR* is integral to the mucoid phenotype (**Figure 6A**). However, restoring mucoidy to Δ*argR* by expressing *rmpD* under its own promoter indicates that RmpD regulates mucoidy downstream of ArgR. This strengthens our model that ArgR regulates mucoidy via the *rmp* promoter.

In the lungs, *K. pneumoniae* faces a complex immune environment where macrophages play a crucial role in pathogen clearance.^16,43,44^ Therefore, we examined how ArgR-dependent mucoidy regulation impacts bacterial interactions with host immune cells, specifically immortalized murine bone marrow-derived macrophages (iBMDMs). We acknowledge that iBMDMs do not fully recapitulate primary alveolar macrophages (AMs). AMs are the first immune cells in the lung to encounter pathogens like *K. pneumoniae* during pulmonary infections, and they are specifically adapted to handle the unique microenvironment of the lung, including interactions with inhaled bacteria.^44,45^ While BMDMs share many functional similarities with AMs and are a widely accepted model for macrophage activation and phagocytosis, they may not fully replicate the immune dynamics encountered in the lung tissue.^45^ Despite this limitation, we observed that ArgR is required for reducing bacterial association with iBMDMs (**Figure 6B**), suggesting that positive regulation of mucoidy by ArgR may facilitate immune evasion by reducing CPS chain length diversity. We speculate that ArgR-mediated mucoidy regulation is a key strategy *K. pneumoniae* uses to evade host immunity, particularly during pulmonary infections. Our data agree with several previous reports, which include that low mucoidy and Δ*rmpD* (resulting in a low mucoid phenotype) increased macrophage association, Δ*argR* increased *Galleria mellonella* survival, and *argR::kan* was significantly out-competed in pneumonia TnSeq and bacteremia TnSeq experiments.^10,11,16,17,40^ Combined, these data suggest that arginine may be a key host cue used by hvKp to optimize *in vivo* fitness and pathogenicity via ArgR-dependent regulation.

Lung infections, particularly pneumonia, are a significant cause of morbidity and mortality worldwide, with *K. pneumoniae* being a major etiological agent.^43,46^ Therefore, to further evaluate the role of ArgR in bacterial fitness, we performed competitive infections between wildtype carrying an empty vector (P_ev_) and Δ*argR* P_ev_ in a murine model of pneumonia. The Δ*argR* P_ev_ mutant exhibited a 2-log decrease in lung colonization (*P* = 0.0008) that was overcome when Δ*argR* P*_argR_* was competed against wildtype P_ev_ (1-log increase, *P =* 0.0078) (**Figure 6C**). These data confirm that ArgR plays an essential role in maintaining bacterial survival and replication in lung tissue, and validate prior TnSeq and insect infection data.^11,16,17,40^ Although Δ*argR* P_ev_ showed a slight trend towards reduced competitive fitness in the liver and spleen, this did not reach statistical significance and was not complemented by Δ*argR* P*_argR_*. The 2-log defect in Δ*argR* primary site colonization (*i.e.* lung) likely confounds our ability to evaluate its role in dissemination to secondary sites. In addition, ArgR also coordinates arginine import and metabolism, host arginine levels fluctuates due to immune responses and inflammation.^47,22,23^ Therefore, the rapid clearance observed in the Δ*argR* mutant may be due to its involvement in these metabolic processes, in addition to its role in mucoidy regulation. Nonetheless, these results demonstrate that ArgR has a critical role in lung colonization, an environment where *K. pneumoniae* encounters a range of immune responses and potentially varying levels of arginine.

Importantly, our results also suggest that arginine regulates mucoidy in multiple hypervirulent strains (**Figure 7A**). The consistent loss of mucoidy in the absence of arginine in two hvKp laboratory strains and three hvKp clinical isolates underscores the likely role arginine abundance plays in regulating hvKp virulence. Finally, we identified a conserved ARG box in the *rmp* promoter (**Figure 7B**). To confirm if the ARG box is required for ArgR to activate P*_rmp_* in KPPR1, we mutated eight conserved ARG box nucleotides in the P*_rmp_* region. P*_rmp_* activity was significantly reduced in all media conditions compared to the wild-type P*_rmp_* vector (**Figure 7C**). This confirms that the ARG box is essential for arginine-dependent regulation of mucoidy. Combined, these data suggest that in the presence of arginine, ArgR directly binds the P*_rmp_* ARG box to positively regulate *rmpD* and increase mucoidy. This presents a promising avenue to further understand how arginine availability regulates hvKp virulence. Further understanding ArgR-P*_rmp_* binding could offer insights for developing new therapies targeting virulence factor regulation, potentially reducing the pathogenicity of hypervirulent *K. pneumoniae*.

Based on data presented here, we propose the following working model (**Figure 8**). Upon bacterial uptake of arginine into the cell, ArgR binds arginine to become activated.^48–50^ In addition to other regulatory targets, the activated ArgR binds to the predicted P*_rmp_* ARG box and activates the *rmp* promoter. This activity increases *rmpD* transcription and RmpD then interacts with Wzc to modulate CPS chain diversity.^11,12^ In the context of bacterial pneumonia, we propose that arginine availability and therefore, ArgR regulation of mucoidy is a method employed by hvKp to modulate the bacterial cell surface and evade the host immune system in the lungs. Here, we demonstrate that ArgR blocks macrophage association and increases bacterial fitness in the lung.

To the best of our knowledge, this study presents the first mechanistic evidence for how ArgR regulates *K. pneumoniae* mucoidy. Our data add to existing literature, which reports that mucoidy is important for avoiding uptake by macrophages.^10,11^ However, further investigation is warranted to understand where *K. pneumoniae* encounters varying arginine concentrations during colonization, invasion and infection, and how arginine-induced changes to *K. pneumoniae* cell surface properties shape the course of pathogenesis. Here, we have noted that glycerol also suppresses mucoidy, and other studies have demonstrated that mucoidy is suppressed when *K. pneumoniae* are cultured in human urine, or in the presence of L-glucose relative to L-fucose.^11,14^ These combined observations lead us to speculate that inputs from multiple signals are integrated in a complex regulatory cascade that fine-tunes CPS chain length diversity and mucoidy in *K. pneumoniae*. We propose that coordinated changes in cell surface properties could block clearance of bacteria by macrophages, thereby increasing bacterial survival in primary host tissues and, potentially, increasing persistence and dissemination to secondary sites. \ Conversely, we speculate that the ability to switch to a non-mucoid phenotype in low-arginine environments (*e.g.* urine) may promote adherence and persistence, without requiring the bacteria to lose capsule production entirely as reported in ST258 lineages.^52^ Fully dissecting the complex regulatory network that optimizes hvKp fitness warrants further exploration and it may identify intervention points that could be targeted by anti-virulence therapies.

In summary, this study reveals the crucial role of arginine as a key signal that regulates mucoidy in *K. pneumoniae* and highlights the ability of this species to fine-tune the cell-surface polysaccharides presented on the cell in response to environmental cues. Based on these data, we posit that *K. pneumoniae* is a ‘capsule expert’ with the ability to fine-tune capsule properties which may optimize its niche-specific fitness. Here, we demonstrated that arginine is essential for enhancing mucoidy independently of capsule abundance. Arginine positively regulates mucoidy by increasing *rmp* promoter activity and *rmpD* transcription, which decreases CPS chain length diversity. Our results indicate that the regulatory mechanisms governing the influence of arginine on mucoidy is consistent across multiple hvKp strains, underscoring its potential significance as a key driver of *K. pneumoniae* hypervirulence. Our findings also suggest that arginine signaling via ArgR may facilitate immune evasion and fitness in the lungs, expanding our understanding of factors shaping *K. pneumoniae* interactions with host immune cells. Finally, we identified a conserved ARG box within the *rmp* operon promoter. It is likely other nutrients or metabolic intermediates exert regulatory control over the *rmp* locus.

Ultimately, this work underscores the complex interactions between nutrient availability and bacterial virulence gene expression. Specific to hypervirulent *K. pneumoniae,* these results reveal that the mucoid phenotype is dynamic and regulated in response to environmental cues. That is, hypervirulent *K. pneumoniae* do not exist in a persistently hypermucoid state. Gaining insights into the regulatory networks that activate virulence factors *in vivo* is crucial for developing new targets that could reduce the pathogenicity of hypervirulent *K. pneumoniae*.

## ACKNOWLEDGMENTS

The following reagent was obtained through BEI Resources, NIAID, NIH: Macrophage Cell Line Derived from Wild-Type Mice, NR-9456. We thank Drs. Neal Hammer and Paige Kies for guidance on supplementing amino acids into a defined medium. We thank Drs. Matthew Parsek and Xuhui Zheng for the pBBR1 plasmid with mScarlet. We thank members of the Mike lab and Department of Medical Microbiology and Immunology at the University of Toledo for critical feedback and for sharing technical expertise and resources, especially Emily Kinney, Krista Pettee, Drs. R. Mark Wooten, Jason Huntley, Jennifer Hill, and Robert Blumenthal.

Research reported in this publication was supported by the University of Toledo College of Medicine and Life Sciences, American Heart Association 23CDA1056712 (L.A.M.), and K22 AI145849 (L.A.M.), R35 GM150588 (L.A.M.), and K99A1175481 (C.L.H.) from the National Institutes of Health. The content is solely the responsibility of the authors and does not necessarily represent the official views of the American Heart Association or the National Institutes of Health.

## MATERIALS AND METHODS

### Bacterial strains and culture conditions

All the primers, bacterial strains, and plasmids described in these studies are detailed in **Tables S1** and **S2.**^30,53,54^ Bacteria were cultured in either lysogeny broth (LB) (5 g/L yeast extract, 10 g/L tryptone, 0.5 g/L NaCl) or low-iron minimal M9 base medium (6 g/L disodium hydrogen phosphate, 3 g/L monopotassium phosphate, 0.5 g/L sodium chloride, 1 g/L ammonium chloride, 14.7 g/L calcium chloride, 120.3 g/L magnesium sulfate) with the reported carbon sources (**Table S3**) at 200 rpm and 37 °C, unless otherwise noted.^19^ Solid medium was prepared by adding 20 g/L bacto-agar to LB medium prior to autoclaving. When appropriate, antibiotics were added at the following concentrations: kanamycin (25 µg/mL), chloramphenicol (20 µg/mL *E. coli* or 80 µg/mL *K. pneumoniae*), gentamicin (10 µg/mL). *K. pneumoniae* KPPR1 mutant strains were constructed using the λ Red recombinase system, as previously described.^19,20^

### Sedimentation Assay

Hypermucoviscosity was quantified using a sedimentation assay as previously described.^11,26^ If experimental conditions yielded an OD_600_ > 0.5, then 0.5 OD_600_ unit of bacterial culture was pelleted in a 2 mL microcentrifuge tube at 1,000 x *g* for 5 min. The OD_600_ of the upper supernatant was then quantified. If experimental conditions resulted in an OD_600_ < 0.5, then 0.5 OD_600_ of bacteria was transferred to a 2 mL microcentrifuge tube and pelleted at 21,000 x *g* for 15 min as previously described.^11^ All but 50 µL of the sample supernatant was removed, then the bacterial pellet was resuspended to 1 mL with PBS, and sedimentation efficiency was quantified after centrifugation at 1,000 x *g* for 5 min.

### Uronic acid quantification

For cell-associated capsular polysaccharide (CPS) quantification in all media conditions, uronic acid quantification was performed as previously described.^11,26^ In brief, 250 µL of the overnight culture was mixed with either 50 μL 1% Zwittergent 3–14 in 100 mM citric acid buffer, pH 2 for CPS. The mixture was incubated at 50 °C for 20 min and then pelleted at 17,000 × g for 5 min. Then, 100 µL of the supernatant was transferred to 400 µL of ice-cold ethanol and incubated for 20 min on ice to precipitate CPS. Samples were rehydrated in 200 µL of ultra-pure water and then 1.2 mL of 0.0125 M sodium tetraborate in concentrated sulfuric acid was added.

### Screening concentrations of arginine and phenylalanine

To identify concentrations of arginine and/or phenylalanine sufficient to restore mucoidy, we cultured KPPR1 in a 96-well plate that had wells filled with 100 µL of low-iron M9 medium supplemented with 0.2% glycerol and had increasing concentrations of phenylalanine and/or arginine. Screening of the concentrations was done as previously described with some adaptations.^29^ To prevent evaporation, the plates were wrapped with plastic wrap and incubated with shaking (200 rpm) at 37°C for 18–20 hours. The sedimentation assay was adapted to a microplate format as follows: Plates were vortexed on low to resuspend pellets, and the total OD_600_ was recorded. Following centrifugation at 1,000 x *g* for 5 minutes, the upper 50 µL of supernatant was transferred to a new microplate containing 50 µL of PBS to measure the OD_600_.

### Growth curves

Bacterial strains were cultured at 200 rpm overnight in triplicate in 3 mL of the relevant medium in aerated culture tubes at 37 °C. The culture OD_600_ was measured and then normalized to an OD_600_ of 0.001 in the reported medium. Subsequently, 100 µL of each back-diluted culture was aliquoted into a microplate with parafilm-wrapped edges to mitigate evaporation-related edge effects. A plate reader (EPOCH2-SN, Agilent) recorded the OD_600_ every 15 min for 16 h. The microplates were incubated at 37 °C with continuous, orbital shaking [282 cpm (3 mm)].

### Molecular cloning and transformation

The oligonucleotides and plasmids employed in this study can be found in Tables S1 and S2. For generating knockouts, λ Red recombineering adapted to *K. pneumoniae*, as previously described, was employed.^11,27–29^ To construct the plasmids, the pACYC184 or pBBR1MCS5-P_J23100_-*mScarlet* backbone and specific gene fragments were PCR amplified, gel-purified (Monarch, NEB), and assembled with NEBuilder HiFi DNA Assembly mix (NEB) for 1 h at 50°C.^55^ The resulting ligated products were transformed into TOP10 *E. coli* through heat-shock. Plasmid construction was confirmed via sequencing. Electroporation of vectors into TOP10 *E. coli* or relevant *K. pneumoniae* strains was performed as previously described.^28^

### Whole-genome sequencing and analysis

Genomic DNA was purified from overnight cultures by pelleting 1 mL of bacterial culture at 15,000 × *g* for 15 min at 4 °C. Samples were prepared according to the directions from the Wizard HMW DNA Extraction kit for Gram-negative Bacteria (Promega).^56^ The genomic DNA was rehydrated in DNA Rehydration Solution and Illumina sequenced (SeqCoast). Sequence variants were detected using the variation analysis pipeline on BV-BRC with the *K. pneumoniae subsp. pneumoniae* ATCC 43816 (Taxonomy ID: 1308539) as the reference genome.^26^

### Western blot

Western blots were performed as previously described.^11^ Whole cell lysates were prepared from 27 mL of overnight culture. The bacterial culture was pelleted at 21,000 x *g* for 15 min at 4 °C and samples were kept on ice. Bacterial pellets were resuspended in 500 µL of lysis buffer; lysis buffer was prepared fresh as follows: one cOmplete Mini, EDTA-free Protease Inhibitor Cocktail tablet (Roche) and one PhosSTOP tablet (Roche) dissolved in 7 mL BugBuster (MilliporeSigma). Each sample was sonicated in two, 5 s pulses with 50% amplitude and 5 s rest between each pulse. Lysed samples were treated with nuclease for 1 h to overnight at 37°C, 200 rpm. The nuclease was prepared fresh (4.9 mg ribonuclease A [Worthington] and 1 mg deoxyribonuclease [Worthington] in 960 µL of 1 M Tris pH 7.5, 20 µL of 1 M CaCl_2_, and 20 µL 1 M MgCl), and 24 µL was added to 500 µL of lysate. Protein concentration was quantified with a BCA assay (Pierce). Samples were prepared at 2 µg/µL in 1× SDS loading buffer and boiled at 70 °C for 3.5 min. 20 µg of protein was resolved on 12% SDS-PAGE gels and transferred to a nitrocellulose membrane then blocked overnight with 5% BSA in TBS. The blot was probed with 1:2,500 mouse anti-phosphotyrosine PY20 (Abcam), then washed with TBS-T, and probed with 1:5,000 goat anti-mouse IgG (H + L)-HRP (secondary) (SouthernBiotech). Blots were developed with ECL Western Blotting Substrate (Pierce) and imaged on a G:Box Imager (Syngene).^11^

### Blot stripping and re-probing

Bound antibodies were stripped off the nitrocellulose membrane with fresh, mild stripping buffer for 10 min, twice. The stripping buffer was prepared at 100 mL by adding 1.5 g glycine, 0.1 g SDS, and 1 mL Tween-20 in water with the final pH adjusted to 2.2. Blots were washed twice with PBS (10 min) and then twice with TBS-T (5 min). Stripped blots were then blocked overnight with 5% BSA in TBS overnight and then re-probed with 1:2,500 anti-GAPDH loading control (primary) antibody (Invitrogen) and 1:5,000 goat anti-mouse IgG (H + L)-HRP (secondary) (SouthernBiotech). Blots were developed as previously described.^11^

### Western blot quantification

Quantification of Western blots was conducted using ImageJ version 1.53 K for Windows. In short, the measurement involved selecting equal areas of each phosphotyrosine-Wzc band and GAPDH band, followed by subtracting the lane background from the obtained values. The background-subtracted value for each band was normalized relative to its corresponding background-subtracted GAPDH value. The resulting ratio of the normalized values for each bacterial strain and media condition was compared to wild type was plotted. The quantification presented in the graphs represents the average of at least three independent replicates.

### Fluorescence reporter assay

Bacterial strains with the fluorescent reporter plasmid were cultured at 200 rpm overnight at 37°C in triplicate in 3 mL of the reported medium in aerated culture tubes. The next day, 1 mL of overnight culture was centrifuged at 21,000 x *g* for 10 min. The supernatant was discarded, and the pellet was washed twice with 1,000 µL of 1x sterile PBS, then centrifuged at 21,000 x *g* for 10 min. The pellet was resuspended in 1 mL of PBS by pipetting. To a 96-well microplate, 300 µL of the undiluted sample was loaded. A blank of 300 µL 1x PBS was also included. The OD_600_ of the samples and dasherGFP abundance (485 ex/528 em) were then measured using a Cytation 5 (Biotek). The transcriptional activity of the *rmp* promoter was calculated by dividing the dasherGFP fluorescence intensity by the OD_600_ for each sample.

### RNA isolation and quantitative RT-PCR

qRT-PCR was performed as previously described with some modifications.^11^ In brief, bacteria cultured overnight in low-iron M9 minimal medium with the appropriate amino acids and 0.2% glycerol were diluted 1:100 into the respective low-iron M9 minimal medium culture. The bacteria were cultured with aeration at 37 °C, 200 rpm for 6 h. Then, approximately 1 × 10^9^ CFU of bacteria were mixed at a 2:1 (vol:vol) ratio of RNAProtect (Qiagen) and incubated at room temperature for 5 min. Samples were then pelleted at 5,000 × *g* for 10 min and the supernatant was decanted. RNA was purified using the RNeasy mini-prep kit (Epoch) after adding 100 µL of lysozyme (15 mg/ml in TE buffer) and 10 µL of proteinase K (20 mg/mL) treatment for 10 min at room temperature. RNA was isolated using an RNA Mini Prep Kit (Epoch) and eluted with 30 µL of RNase-free water and samples were stored at -20 °C. Genomic DNA was removed using ezDNAse (ThermoFisher) and then cDNA synthesis was performed with SuperScript IV Reverse Transcriptase (Invitrogen) on an equal amount of RNA for each sample, roughly 1 µg total. The resulting cDNA was diluted 1:50 in water and used as template for quantitative real-time PCR (qRT-PCR) in a QuantStudio 3 PCR system (Applied Biosystem) with SYBRGreen PowerUp reagent (Invitrogen). Primers for amplifying internal fragments of the *rmpD* and *gap2* genes are listed in **Table S1**, with the *gap2* transcript serving as an internal control.^11^ The relative fold change was calculated using the comparative threshold cycle (C_T_) method as previously described.^57^

### CPS chain length visualization

CPS chain length visualization was performed as previously described with some modifications.^11,26^ In brief, 1.5 OD_600_ of bacteria cultured overnight in low-iron M9 minimal medium with the appropriate amino acids and 0.2% glycerol were transferred to a microcentrifuge tube and pelleted at 21,000 x *g* for 15 min. The bacterial pellet was washed with 1 mL of sterile 1x PBS and centrifuged again at 21,000 x *g* for 15 min and then discarded 750 µL of the supernatant without disrupting the pellet. The pelleted cells were resuspended with the remaining volume of 250 µL PBS, and then 50 µL of 3–14 Zwittergent was added and incubated at 50 °C for 20 min. Samples were then centrifuged at 17,000 × *g* for 5 min and 100 µL of upper supernatant were precipitated in 400 µL of ice-cold ethanol for 20 min. Samples were then centrifuged at 17,000 × *g* for 5 min at 4 °C and the supernatant was discarded. The pelleted cells were then resuspended with 200 µL of ultra-pure water and placed at 37 °C for 30 min. To prepare the samples for SDS-PAGE, 75 µL of the sample was added to 25 µL of 4x SDS loading buffer. Twenty microliters of prepared sample were loaded on a 4-15% Mini-PROTEAN TGX stain-free pre-cast gel (Bio-Rad) with 20 µL of Precision Plus All Blue standard (Bio-Rad). The polysaccharides were electrophoresed for 4.5 h at 300 V on ice at 4 °C. After electrophoresis, the gel was washed in 200 mL ultrapure water for 10 min, a total of five times. It was then stained in 0.1% alcian blue stain (0.1% wt/vol ThermoFisher Alcian Blue 8 GX in stain base solution) for 1 h with rocking. Stain base solution was prepared as 40% ethanol and 60% 20 mM sodium acetate, pH 4.75. After staining, the gel was de-stained in stain base solution overnight with rocking and then stained using Pierce Silver Stain Kit (ThermoFisher) and imaged on a Syngene G:box using the visible protein setting.

### CPS chain length quantification

Quantification of the stained gels was conducted using ImageJ version 1.53 K for Windows. In brief, equal areas of the entire lane were selected and then the lane background was subtracted from the obtained values. The background-subtracted value for each identified CPS chain type for each bacterial strain and media condition was plotted as area under the curve. The quantification presented in the graphs represents the average of at least three independent replicates.

### Macrophage association assay

Macrophage association assays were performed as previously described with the following modifications.^29^ Immortalized macrophage cells derived from wild-type mice (BEI Resources #NR-9456) were maintained in DMEM medium with L-glutamine, 4.5 g/L glucose and sodium pyruvate (Corning) supplemented with 10% heat-inactivated fetal calf serum (Corning), 100 U/mL penicillin, and 100 µg/mL streptomycin in an atmosphere of 5% CO_2_. Cells were checked for confluency (∼3 × 10^5^ cells/well) in 24-well tissue culture dishes and were then washed with 1 mL of DPBS. Then, 1 mL of 3 × 10^6^ CFU/mL bacteria (MOI 10) in unmodified DMEM medium was added to each well. The 24-well tissue culture plate with samples were spun at 500 rpm (54 × *g*) for 5 min to bring the bacteria in contact with the macrophages and then incubated at 37°C, 5% CO_2_ for 2 h. Samples were washed three times with sterile 1x PBS and lysed with 1 mL of 0.2% Triton-X100 in PBS for 5 min on a rocker. Input and cell-associated bacterial counts were determined by serial dilution and CFU enumeration on LB agar.

### Murine pneumonia model

The murine pneumoniae model was performed as previously described.^58^ In brief, this study was performed using 7-10 week-old mice (Jackson Laboratory, Bar Harbor, ME) adhering to humane animal handling recommendations and approved by the University of Michigan Institutional Animal Care and Use Committee (protocol: PRO00011097). Overnight LB cultures with the appropriate antibiotics *of K. pneumoniae* were centrifuged, resuspended, and adjusted to the proper concentration in PBS. Mice were anesthetized using isoflurane and a total 1x10^6^ CFU *K. pneumoniae* in a 50 µL volume (1:1 ratio of WT:mutant) was administered retropharyngeally. Twenty-four hours postinfection, mice were euthanized by carbon dioxide asphyxiation prior to collection of lung, spleen, and liver. Organs were homogenized in sterile PBS, serially diluted and CFU enumerated on LB agar with appropriate antibiotics. The competitive index was defined as CFU from (mutant output/wild-type output)/(mutant input/wild-type input).

### Genetic characterization of hvKp clinical isolates and ARG box alignment

Raw sequence reads of Kp4289 (SRA accession: SRR19095630), Kp4585 (SRA accession: SRR19095615), and Kp6557 (SRA accession: SRR18982352) from the NCBI database were assembled using PATRIC (https://www.bv-brc.org).^30^ Assembled sequences, including NTUH-K2044, were uploaded to Pathogenwatch (https://pathogen.watch/) for species identification and identification of the *rmp* operon.^31–37^ The *rmp* promoter of each of the above SRA-deposited genomes and NTUH-K2044 was identified by BLAST using the promoter region from KPPR1. The promoters were aligned to a previously reported ARG box consensus sequence using CLUSTALW.^39,59^

### Statistics

All replicates represent biological replicates, except for qPCR which was technical replicates and were replicated at least three times. All statistical analyses were computed in Prism 10 (GraphPad Software, La Jolla, CA, USA). For experiments comparing multiple groups on two independent factors, significance was calculated using two-way ANOVA with either a Tukey or Bonferroni posttest to compare specific groups. For experiments comparing three or more groups on one independent factor, significance was calculated one-way ANOVA either a Dunnett or Tukey posttest. A student t-test was applied when comparing two groups or a single group to a hypothetical value of 1.00. Results were considered significant if the P value was less than or equal to 0.05. All data presented are the mean, and error bars represent the standard error of the mean.

---

**Supplementary Figure 1.**
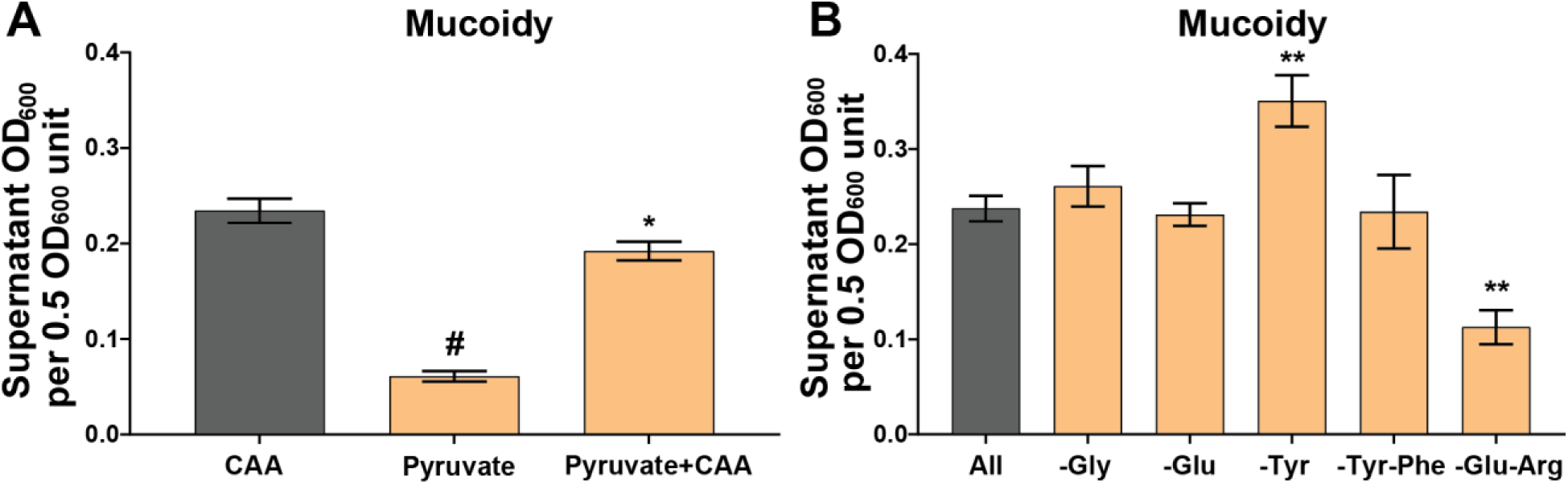
Pyruvate suppresses mucoidy and amino acids not required for increased mucoidy. KPPR1 was cultured overnight in (**A**) low-iron M9 minimal medium with either 1% casamino acids (CAA), 20 mM sodium pyruvate, or both as nutrient sources. Additionally, KPPR1 was cultured overnight in (**B**) low-iron M9 minimal medium with the following amino acids absent: glycine (-Gly), glutamate(-Glu), tyrosine(-Tyr), tyrosine and phenylalanine(-Tyr-Phe), or arginine and glutamate (-Arg-Glu). (**A, B**) Mucoidy was determined for all the different media compositions by quantifying the supernatant OD600 after sedimenting 0.5 OD600 unit of culture at 1,000 x *g* for 5 min. In this supplementary figure, data reused from Figure 1A and Figure 2C is represented by dark grey bars to facilitate comparisons. Data presented are the mean, and error bars represent the standard error of the mean. Statistical significance was determined using a one-way ANOVA and Dunnett’s post-test relative to (**A**) CAA and (**B**) M9+All. * p< 0.05; ** p < 0.01; # p < 0.0001. Experiments were performed ≥3 independent times, in triplicate.

**Supplementary Figure 2.**
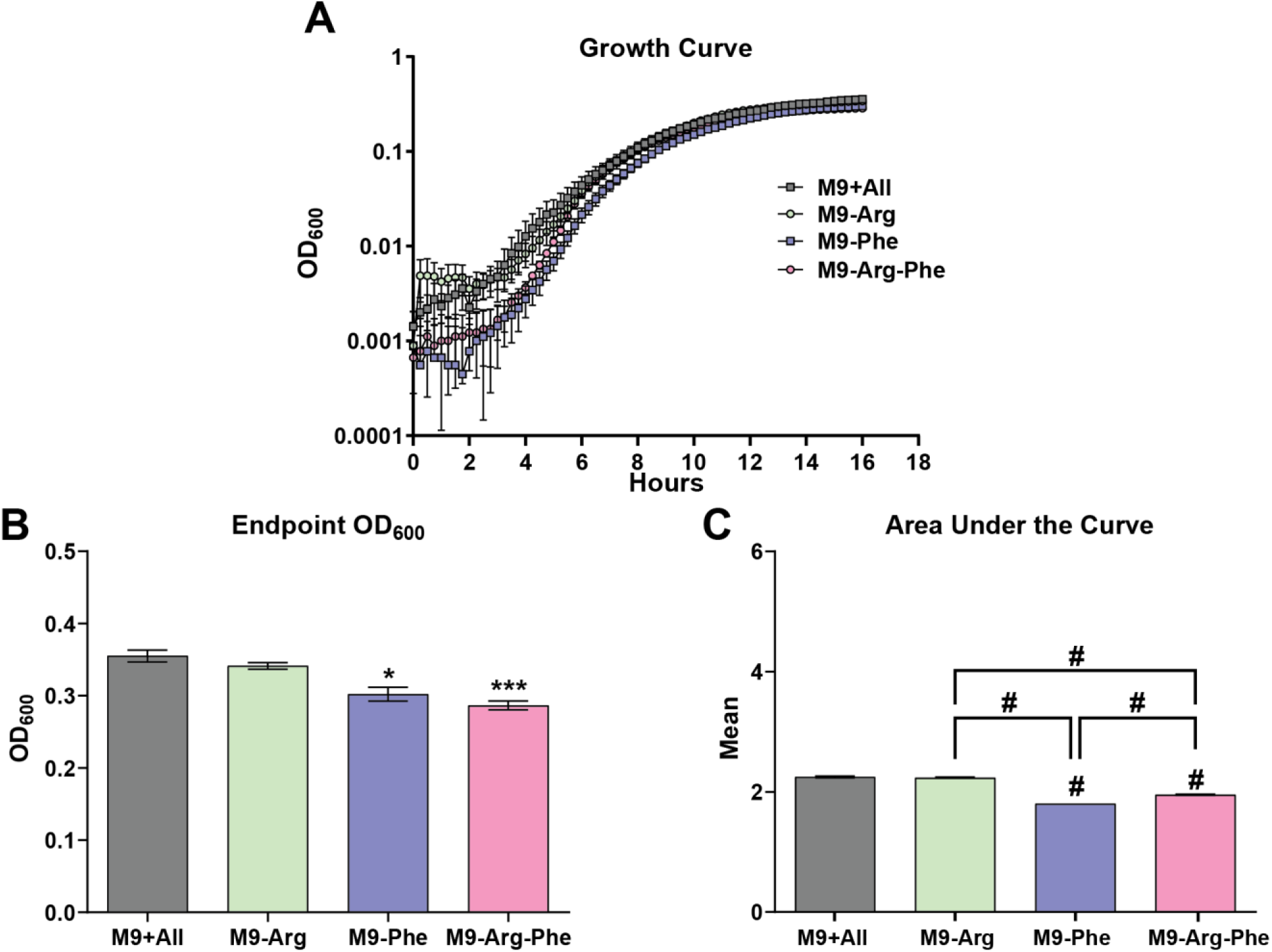
Absence of arginine does not affect KPPR1 growth. Wild type *K. pneumoniae* strain, KPPR1, was cultured overnight in low-iron M9 minimal medium with either 18 amino acids (M9+All), arginine absent (M9-Arg), phenylalanine absent (M9-Phe), or both arginine and phenylalanine absent (M9-Arg-Phe), then back-diluted to OD_600_ 0.0001 in the respective media. Cultures were incubated at 37°C with continuous shaking. (**A**) OD_600_ measurements were collected every 15 min. Data presented are the mean, and error bars represent the standard error of the mean. To evaluate growth in the four culture conditions, (**B**) endpoint OD_600_ and (**C**) area under the curve was assessed for total growth and cumulative growth, respectively. The statistics displayed above the data bars represent values relative to the LB medium. The additional statistics connected by lines serve as direct comparisons to highlight differences between M9 media conditions. Statistical significance was determined using a (**B**) Kruskal-Wallis with Dunn’s post-test and (**C**) one-way ANOVA with Tukey’s post-test. The statistics displayed above the data bars represent values relative to the M9+All. * p < 0.05; *** p < 0.001; # p < 0.0001. Experiments were performed ≥3 independent times, in triplicate.

**Supplementary Figure 3.**
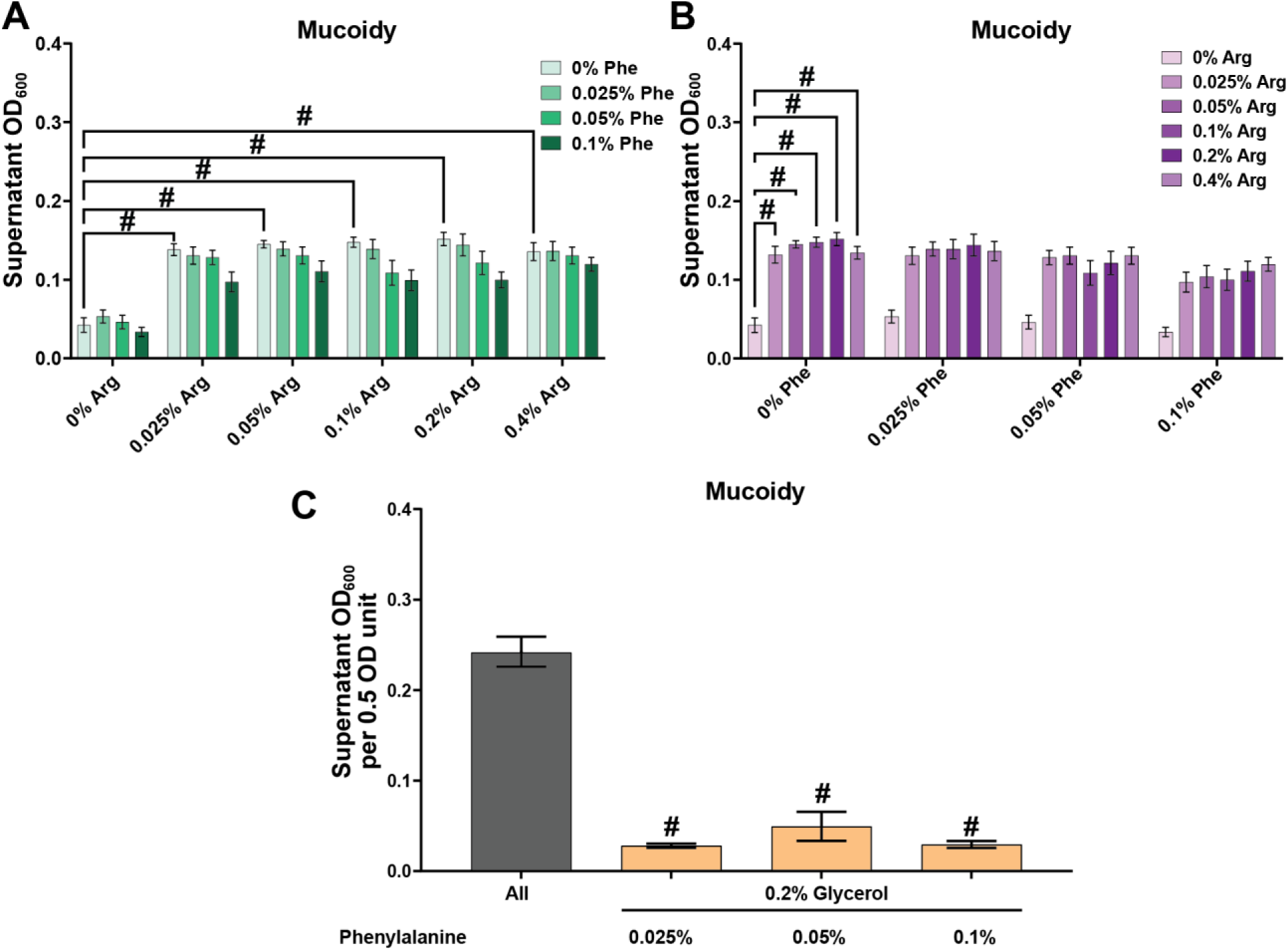
Phenylalanine is not able to restore mucoidy alone. KPPR1 was cultured (**A,B**) 96 well plate that had wells filled with 100 µL of low-iron M9 medium supplemented with 0.2% glycerol and had increasing concentrations of phenylalanine and arginine and (**C**) cultured in 16 mL tubes with all 18 amino acids (All) or in only phenylalanine (0.025%, 0.05%, and 0.1%) and 0.2% glycerol. Both panels **A** and **B** represent the same data, but both are presented for clarity with concentrations of arginine on the x-axis or concentrations of phenylalanine, respectively. (**A,B**) Mucoidy was measured by resuspending the pellets in the 96-well plate by gentle vortexing and the total OD_600_ was measured. The 96 well plate was centrifuged at 1,000 x *g* for 5 minutes and the upper 50 µL OD_600_ of the supernatant was measured. **(C)** Mucoidy was determined for the increasing concentrations of phenylalanine and 0.2% glycerol by quantifying the supernatant OD600 after sedimenting 0.5 OD600 unit of culture at 1,000 x *g* for 5 min. In this supplementary figure, data reused from Figure 2C is represented by a dark grey bar to facilitate comparisons. Data presented are the mean, and error bars represent the standard error of the mean. Statistical significance was determined by (**A,B**) using a two-way ANOVA and Tukey’s post-test and (**B**) comparing the phenylalanine concentrations to the M9+All condition using a one-way ANOVA and Dunnett’s post-test. # p < 0.0001. Experiments were performed ≥3 independent times, in triplicate.

**Supplementary Figure 4.**
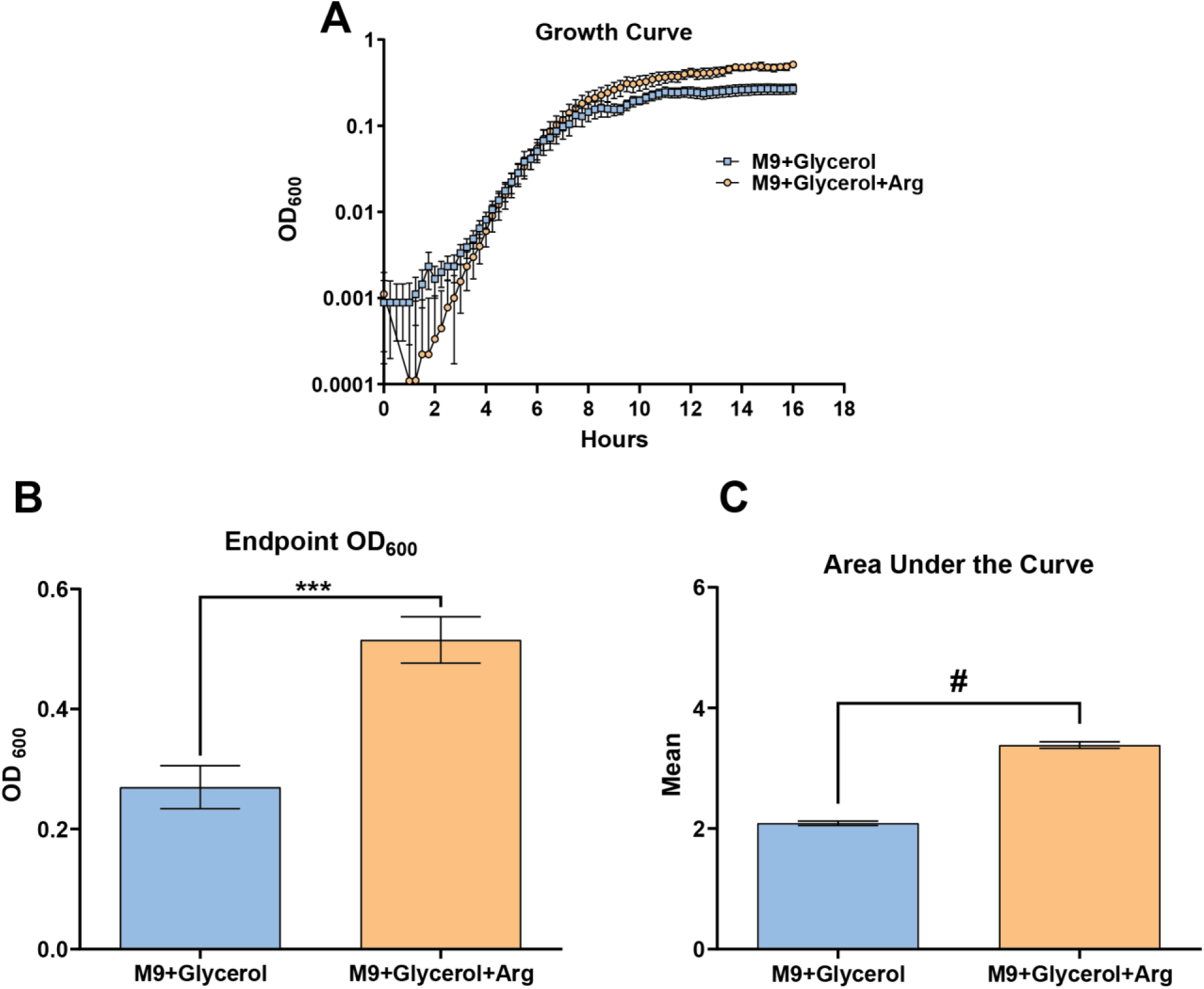
M9+Glycerol yields less KPPR1 growth. Wild type *K. pneumoniae* strain, KPPR1, was cultured overnight in low-iron M9 minimal medium with either glycerol (M9+Glycerol) or glycerol and arginine (M9+Glycerol+Arg), then back-diluted to OD_600_ 0.0001 in the respective media. Cultures were incubated at 37°C with continuous shaking. (**A**) OD_600_ measurements were collected every 15 min. Data presented are the mean, and error bars represent the standard error of the mean. To evaluate growth in the 2 culture conditions, (**B**) endpoint OD_600_ and (**C**) area under the curve was assessed for total growth and cumulative growth, respectively. Statistical significance was determined using an unpaired Student’s t-test. *** p < 0.001; # p < 0.0001. Experiments were performed >3 independent times, in triplicate.

**Supplementary Figure 5.**
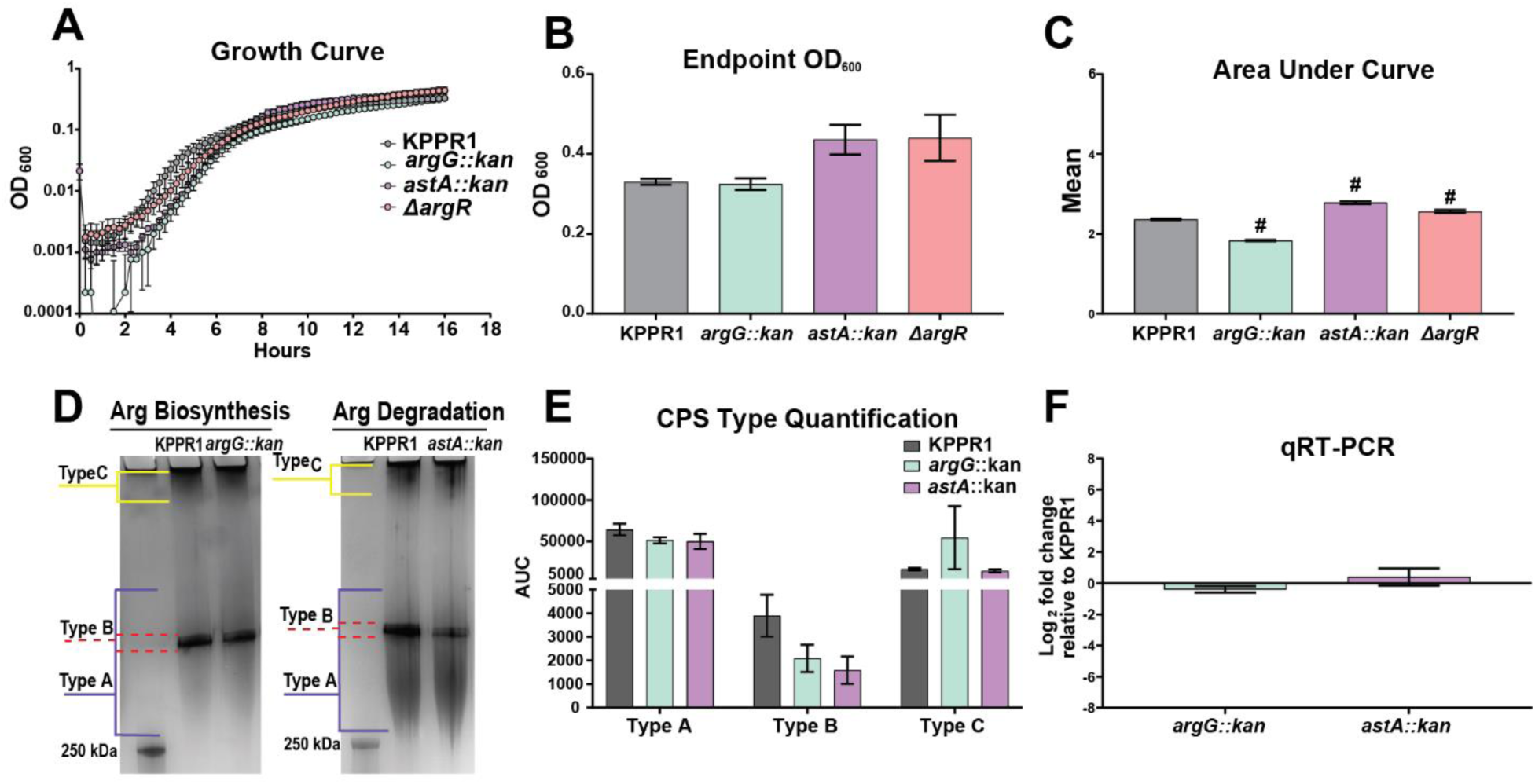
All three of the mutants affect cumulative growth, but not overall growth, and AstA and ArgG do not impact capsule chain diversity or *rmpD* transcripts. Wild type *K. pneumoniae* strain, KPPR1, *argG::kan*, *astA::kan*, and Δ*argR* mutant were cultured overnight in low-iron M9 minimal medium with 18 amino acids (M9+All), then back-diluted to OD_600_ 0.0001 in the medium. Cultures were incubated at 37°C with continuous shaking. (**A**) OD_600_ measurements were collected every 15 min. Data presented are the mean, and error bars represent the standard error of the mean. To evaluate growth in the two culture conditions, (**B**), endpoint OD_600_ and (**C**) area under the curve was assessed for total growth and cumulative growth, respectively. The two mutant strains, *argG*::kan and *astA*::kan, were cultured in M9+All medium overnight. (**D**) Purified capsular polysaccharides (CPS) were separated on a 4-15% SDS-PAGE gel and stained with alcian blue and silver stain. (**E**) Quantification of each capsule chain type was done using ImageJ. Data presented are the mean, and error bars represent the standard error of the mean. (**F**) The relative abundance of *rmpD* for each of the mutants was determined by qRT-PCR and normalized to *gap2* transcript abundance. Relative *rmpD* transcript levels were measured for each mutant and then compared to the wildtype, KPPR1. Representative images are shown in panel **D**. Statistical significance was determined using a one-way ANOVA with Dunnett’s post-test. # p < 0.0001. Experiments were performed >3 independent times, in triplicate.

**Supplementary Figure 6.**
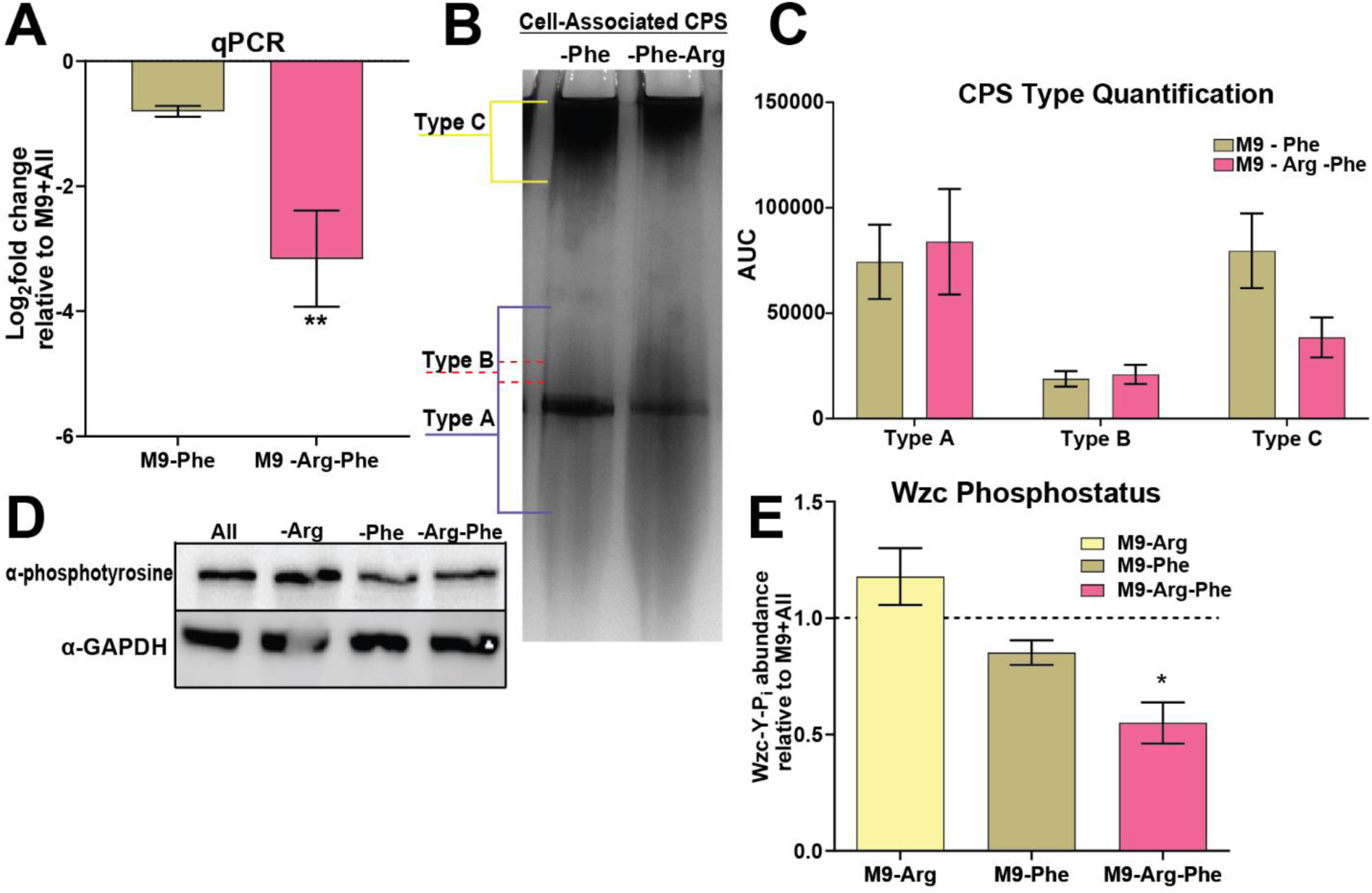
Phenylalanine alone does not affect capsule chain diversity or *rmpD* transcription, but its absence with arginine alters Wzc phospho-status and *rmpD* transcription. KPPR1 was cultured in M9 minimal medium with all amino acids except phenylalanine (M9-Phe) or phenylalanine and arginine (M9-Arg-Phe) to mid-log and RNA was isolated. (**A**) The relative abundance of *rmpD* in the different conditions was determined by qRT-PCR and normalized to *gap2* transcript abundance. Relative *rmpD* transcript levels were measured in the two conditions and then compared to M9+All. KPPR1 was cultured in the two conditions as indicated above each lane. (**B**) Capsular polysaccharides (CPS) were separated on a 4-15% SDS-PAGE gel and stained with alcian blue and silver stain. (**C**) Quantification of each capsule chain type was done using ImageJ. Whole-cell lysates from KPPR1 in the M9 minimal medium with 18 amino acids (M9+All), in the absence of arginine (M9-Arg), with glycerol only (M9+Glycerol), or glycerol and arginine (M9+Glycerol+Arg) were (**D**) probed with anti-phosphotyrosine antibody and anti-GAPDH antibody in a western blot. (**E**) Quantification of Wzc-phosphotyrosine bands from (**D**) using ImageJ, normalized to GAPDH, and relative to M9+All.. Data presented are the mean, and error bars represent the standard error of the mean. Representative images are shown in panels **B** and **D**. Statistical significance was determined in panels **A** and **E,** a one-way ANOVA with a Dunnett’s post-test was used to determine if either group were significantly different than 1.0. In panel **C** a one-way ANOVA with Bonferroni post-test was used to compare the two conditions relative to M9+All, not pictured here. * p<0.05; ** p < 0.01. Experiments were performed ≥3 independent times, in triplicate.

## REFERENCES

1 Ikuta, K. S. et al. Global mortality associated with 33 bacterial pathogens in 2019: a systematic analysis for the Global Burden of Disease Study 2019. The Lancet 400, 2221–2248 (2022).

2 Lan, P., Jiang, Y., Zhou, J. & Yu, Y. A global perspective on the convergence of hypervirulence and carbapenem resistance in Klebsiella pneumoniae. J Glob Antimicrob Resist 25, 26–34 (2021). 10.1016/j.jgar.2021.02.020

3 Russo, T. A. & Marr, C. M. Hypervirulent Klebsiella pneumoniae. Clin Microbiol Rev 32 (2019). 10.1128/cmr.00001-19

4 Russo, T. A. et al. Identification of Biomarkers for Differentiation of Hypervirulent Klebsiella pneumoniae from Classical K. pneumoniae. J Clin Microbiol 56 (2018). 10.1128/jcm.00776-18

5 Dai, P. & Hu, D. The making of hypervirulent Klebsiella pneumoniae. J Clin Lab Anal 36, e24743 (2022). 10.1002/jcla.24743

6 Chang, D., Sharma, L., Dela Cruz, C. S. & Zhang, D. Clinical Epidemiology, Risk Factors, and Control Strategies of Klebsiella pneumoniae Infection. Frontiers in Microbiology 12 (2021). 10.3389/fmicb.2021.750662

7 Fang, C. T., Yi, W. C., Shun, C. T. & Tsai, S. F. DNA adenine methylation modulates pathogenicity of Klebsiella pneumoniae genotype K1. J Microbiol Immunol Infect 50, 471–477 (2017). 10.1016/j.jmii.2015.08.022

8 Choby, J. E., Howard-Anderson, J. & Weiss, D. S. Hypervirulent Klebsiella pneumoniae – clinical and molecular perspectives. Journal of Internal Medicine 287, 283–300 (2020). 10.1111/joim.13007

9 Piperaki, E. T., Syrogiannopoulos, G. A., Tzouvelekis, L. S. & Daikos, G. L. Klebsiella pneumoniae: Virulence, Biofilm and Antimicrobial Resistance. Pediatr Infect Dis J 36, 1002–1005 (2017). 10.1097/inf.0000000000001675

10 Walker, K. A., Treat, L. P., Sepúlveda, V. E. & Miller, V. L. The Small Protein RmpD Drives Hypermucoviscosity in Klebsiella pneumoniae. mBio 11 (2020). 10.1128/mBio.01750-20

11 Khadka, S., et al. Urine-mediated suppression of Klebsiella pneumoniae mucoidy is counteracted by spontaneous Wzc variants altering capsule chain length. Msphere, e00288–00223 (2023).

12 Ovchinnikova, O. G. et al. Hypermucoviscosity Regulator RmpD Interacts with Wzc and Controls Capsular Polysaccharide Chain Length. mBio 14, e0080023 (2023). 10.1128/mbio.00800-23

13 Le, M. N. et al. Genomic epidemiology and temperature dependency of hypermucoviscous Klebsiella pneumoniae in Japan. Microb Genom 8 (2022). 10.1099/mgen.0.000827

14 Hudson, A. W., Barnes, A. J., Bray, A. S., Ornelles, D. A. & Zafar, M. A. Klebsiella pneumoniae l-Fucose Metabolism Promotes Gastrointestinal Colonization and Modulates Its Virulence Determinants. Infect Immun 90, e0020622 (2022). 10.1128/iai.00206-22

15 Carfrae, L. A. & Brown, E. D. Nutrient stress is a target for new antibiotics. Trends Microbiol 31, 571–585 (2023). 10.1016/j.tim.2023.01.002

16 Bachman, M. A. et al. Genome-Wide Identification of Klebsiella pneumoniae Fitness Genes during Lung Infection. mBio 6, e00775 (2015). 10.1128/mBio.00775-15

17 Holmes, C. L. et al. Klebsiella pneumoniae causes bacteremia using factors that mediate tissue-specific fitness and resistance to oxidative stress. PLoS Pathog 19, e1011233 (2023). 10.1371/journal.ppat.1011233

18 Alteri, C. J. & Mobley, H. L. T. Metabolism and Fitness of Urinary Tract Pathogens. Microbiol Spectr 3 (2015). 10.1128/microbiolspec.MBP-0016-2015

19 Murdoch, C. C. & Skaar, E. P. Nutritional immunity: the battle for nutrient metals at the host-pathogen interface. Nat Rev Microbiol 20, 657–670 (2022). 10.1038/s41579-022-00745-6

20 Gogoi, M., Datey, A., Wilson, K. T. & Chakravortty, D. Dual role of arginine metabolism in establishing pathogenesis. Curr Opin Microbiol 29, 43–48 (2016). 10.1016/j.mib.2015.10.005

21 Reitzer, L. & Zimmern, P. Rapid Growth and Metabolism of Uropathogenic Escherichia coli in Relation to Urine Composition. Clin Microbiol Rev 33 (2019). 10.1128/cmr.00101-19

22 Paulson, N. B. et al. The arginine decarboxylase pathways of host and pathogen interact to impact inflammatory pathways in the lung. PLoS One 9, e111441 (2014). 10.1371/journal.pone.0111441

23 Palmer, K. L., Aye, L. M. & Whiteley, M. Nutritional cues control Pseudomonas aeruginosa multicellular behavior in cystic fibrosis sputum. J Bacteriol 189, 8079–8087 (2007). 10.1128/jb.01138-07

24 Wong Fok Lung, T., et al. Klebsiella pneumoniae induces host metabolic stress that promotes tolerance to pulmonary infection. Cell Metab 34, 761–774.e769 (2022). 10.1016/j.cmet.2022.03.009

25 Lin, C. T. et al. Fur regulation of the capsular polysaccharide biosynthesis and iron-acquisition systems in Klebsiella pneumoniae CG43. Microbiology (Reading) 157, 419–429 (2011). 10.1099/mic.0.044065-0

26 Khadka, S., Ring, B. E., Pariseau, D. A. & Mike, L. A. Characterization of Klebsiella pneumoniae Extracellular Polysaccharides. Curr Protoc 3, e937 (2023). 10.1002/cpz1.937

27 Datsenko, K. A. & Wanner, B. L. One-step inactivation of chromosomal genes in Escherichia coli K-12 using PCR products. Proc Natl Acad Sci U S A 97, 6640–6645 (2000). 10.1073/pnas.120163297

28 Ring, B. E., Khadka, S., Pariseau, D. A. & Mike, L. A. Genetic Manipulation of Klebsiella pneumoniae. Curr Protoc 3, e912 (2023). 10.1002/cpz1.912

29 Mike, L. A. et al. A systematic analysis of hypermucoviscosity and capsule reveals distinct and overlapping genes that impact Klebsiella pneumoniae fitness. PLoS Pathog 17, e1009376 (2021). 10.1371/journal.ppat.1009376

30 Vornhagen, J. et al. Combined comparative genomics and clinical modeling reveals plasmid-encoded genes are independently associated with Klebsiella infection. Nat Commun 13, 4459 (2022). 10.1038/s41467-022-31990-1

31 Bayliss, S. C. et al. The Promise of Whole Genome Pathogen Sequencing for the Molecular Epidemiology of Emerging Aquaculture Pathogens. Frontiers in Microbiology 8 (2017). 10.3389/fmicb.2017.00121

32 Harris, S. R. et al. Public health surveillance of multidrug-resistant clones of Neisseria gonorrhoeae in Europe: a genomic survey. Lancet Infect Dis 18, 758–768 (2018). 10.1016/s1473-3099(18)30225-1

33 Sánchez-Busó, L. et al. Europe-wide expansion and eradication of multidrug-resistant Neisseria gonorrhoeae lineages: a genomic surveillance study. Lancet Microbe 3, e452–e463 (2022). 10.1016/s2666-5247(22)00044-1

34 Gladstone, R. A. et al. Visualizing variation within Global Pneumococcal Sequence Clusters (GPSCs) and country population snapshots to contextualize pneumococcal isolates. Microb Genom 6 (2020). 10.1099/mgen.0.000357

35 Mitchell, C. et al. Globetrotting strangles: the unbridled national and international transmission of Streptococcus equi between horses. Microb Genom 7 (2021). 10.1099/mgen.0.000528

36 Argimón, S. et al. A global resource for genomic predictions of antimicrobial resistance and surveillance of Salmonella Typhi at pathogenwatch. Nat Commun 12, 2879 (2021). 10.1038/s41467-021-23091-2

37 Li, X. et al. Comparing genomic variant identification protocols for Candida auris. Microb Genom 9 (2023). 10.1099/mgen.0.000979

38 Solovyev, V. & Salamov, A. in *In* Metagenomics and its Applications in Agriculture, Biomedicine and Environmental Studies *(Ed.* R.W. Li) 61–78 (Nova Science Publishers, 2011).

39 Ramirez, M. S., Traglia, G. M., Lin, D. L., Tran, T. & Tolmasky, M. E. Plasmid-Mediated Antibiotic Resistance and Virulence in Gram-Negatives: the Klebsiella pneumoniae Paradigm. Microbiol Spectr 2 (2014). 10.1128/microbiolspec.PLAS-0016-2013

40. Dorman, M. J., Feltwell, T., Goulding, D. A., Parkhill, J. & Short, F. L. The Capsule Regulatory Network of Klebsiella pneumoniae Defined by density-TraDISort. mBio, 9, (2018). doi:10.1128/mbio.01863-18

41 Cho, B. K., Federowicz, S., Park, Y. S., Zengler, K. & Palsson, B. Deciphering the transcriptional regulatory logic of amino acid metabolism. Nat Chem Biol 8, 65–71 (2011). 10.1038/nchembio.710

42 Barrientos-Moreno, L., Molina-Henares, M. A., Ramos-González, M. I. & Espinosa-Urgel, M. Role of the Transcriptional Regulator ArgR in the Connection between Arginine Metabolism and c-di-GMP Signaling in Pseudomonas putida. Appl Environ Microbiol 88, e0006422 (2022). 10.1128/aem.00064-22

43 Holmes, C. L., Anderson, M. T., Mobley, H. L. T. & Bachman, M. A. Pathogenesis of Gram-Negative Bacteremia. Clin Microbiol Rev 34 (2021). 10.1128/cmr.00234-20

44 Broug-Holub, E. et al. Alveolar macrophages are required for protective pulmonary defenses in murine Klebsiella pneumonia: elimination of alveolar macrophages increases neutrophil recruitment but decreases bacterial clearance and survival. Infect Immun 65, 1139–1146 (1997). 10.1128/iai.65.4.1139-1146.1997

45 Woods, P. S. et al. Tissue-Resident Alveolar Macrophages Do Not Rely on Glycolysis for LPS-induced Inflammation. Am J Respir Cell Mol Biol 62, 243–255 (2020). 10.1165/rcmb.2019-0244OC

46 Cortés, G., Alvarez, D., Saus, C. & Albertí, S. Role of lung epithelial cells in defense against Klebsiella pneumoniae pneumonia. Infect Immun 70, 1075–1080 (2002). 10.1128/iai.70.3.1075-1080.2002

47 Charlier, D. & Bervoets, I. Regulation of arginine biosynthesis, catabolism and transport in Escherichia coli. Amino Acids 51, 1103–1127 (2019). 10.1007/s00726-019-02757-8

48 Torres Montaguth, O. E., Bervoets, I., Peeters, E. & Charlier, D. Competitive Repression of the artPIQM Operon for Arginine and Ornithine Transport by Arginine Repressor and Leucine-Responsive Regulatory Protein in Escherichia coli. Front Microbiol 10, 1563 (2019). 10.3389/fmicb.2019.01563

49 Cho, S. et al. The architecture of ArgR-DNA complexes at the genome-scale in Escherichia coli. Nucleic Acids Res 43, 3079–3088 (2015). 10.1093/nar/gkv150

50 Cherney, L. T., Cherney, M. M., Garen, C. R., Lu, G. J. & James, M. N. Crystal structure of the arginine repressor protein in complex with the DNA operator from Mycobacterium tuberculosis. J Mol Biol 384, 1330–1340 (2008). 10.1016/j.jmb.2008.10.015

51 Lee, H. et al. Evolution of Klebsiella pneumoniae with mucoid and non-mucoid type colonies within a single patient. Int J Med Microbiol 309, 194–198 (2019). 10.1016/j.ijmm.2019.03.003

52 Ernst, C. M. et al. Adaptive evolution of virulence and persistence in carbapenem-resistant Klebsiella pneumoniae. Nat Med 26, 705–711 (2020). 10.1038/s41591-020-0825-4

53 Broberg, C. A., Wu, W., Cavalcoli, J. D., Miller, V. L. & Bachman, M. A. Complete Genome Sequence of Klebsiella pneumoniae Strain ATCC 43816 KPPR1, a Rifampin-Resistant Mutant Commonly Used in Animal, Genetic, and Molecular Biology Studies. Genome Announc 2 (2014). 10.1128/genomeA.00924-14

54 Wu, K. M. et al. Genome sequencing and comparative analysis of Klebsiella pneumoniae NTUH-K2044, a strain causing liver abscess and meningitis. J Bacteriol 191, 4492–4501 (2009). 10.1128/jb.00315-09

55 Anderson, M. T. et al. Identification of distinct capsule types associated with Serratia marcescens infection isolates. PLoS Pathogens 18 (2022).

56 Pariseau, D. A., Ring, B. E., Khadka, S. & Mike, L. A. Cultivation and Genomic DNA Extraction of Klebsiella pneumoniae. Curr Protoc 4, e932 (2024). 10.1002/cpz1.932

57 Schmittgen, T. D. & Livak, K. J. Analyzing real-time PCR data by the comparative C(T) method. Nat Protoc 3, 1101–1108 (2008). 10.1038/nprot.2008.73

58 Holmes, C. L. et al. The ADP-Heptose Biosynthesis Enzyme GmhB is a Conserved Gram-Negative Bacteremia Fitness Factor. Infect Immun 90, e0022422 (2022). 10.1128/iai.00224-22

59 Thompson, J. D., Higgins, D. G. & Gibson, T. J. CLUSTAL W: improving the sensitivity of progressive multiple sequence alignment through sequence weighting, position-specific gap penalties and weight matrix choice. Nucleic Acids Res 22, 4673–4680 (1994). 10.1093/nar/22.22.4673

